# Cell-type-specific regulation of APOE levels in human neurons by the Alzheimer’s disease risk gene SORL1

**DOI:** 10.1101/2023.02.25.530017

**Authors:** Hyo Lee, Aimee J. Aylward, Richard V. Pearse, Yi-Chen Hsieh, Zachary M. Augur, Courtney R. Benoit, Vicky Chou, Allison Knupp, Cheryl Pan, Srilakshmi Goberdhan, Duc M. Duong, Nicholas T. Seyfried, David A. Bennett, Hans-Ulrich Klein, Philip L. De Jager, Vilas Menon, Jessica E. Young, Tracy L. Young-Pearse

**Affiliations:** Ann Romney Center for Neurologic Diseases, Department of Neurology, Brigham and Women’s Hospital and Harvard Medical School, Boston, MA, USA; Department of Laboratory Medicine and Pathology, University of Washington, Seattle, WA, USA; Department of Biochemistry, Emory School of Medicine, Atlanta, GA, USA; Rush Alzheimer’s Disease Center, Rush University Medical Center, Chicago, IL, USA; Center for Translational and Computational Neuroimmunology, Department of Neurology and the Taub Institute for the Study of Alzheimer’s Disease and the Aging Brain, Columbia University Irving Medical Center, New York, NY, USA

## Abstract

SORL1 is strongly implicated in the pathogenesis of Alzheimer’s disease (AD) through human genetic studies that point to an association of reduced SORL1 levels with higher risk for AD. To interrogate the role(s) of SORL1 in human brain cells, SORL1 null iPSCs were generated, followed by differentiation to neuron, astrocyte, microglia, and endothelial cell fates. Loss of SORL1 led to alterations in both overlapping and distinct pathways across cell types, with the greatest effects in neurons and astrocytes. Intriguingly, SORL1 loss led to a dramatic neuron-specific reduction in APOE levels. Further, analyses of iPSCs derived from a human aging cohort revealed a neuron-specific linear correlation between SORL1 and APOE RNA and protein levels, a finding validated in human post-mortem brain. Pathway analysis implicated intracellular transport pathways and TGF- β/SMAD signaling in the function of SORL1 in neurons. In accord, enhancement of retromer-mediated trafficking and autophagy rescued elevated phospho-tau observed in SORL1 null neurons but did not rescue APOE levels, suggesting that these phenotypes are separable. Stimulation and inhibition of SMAD signaling modulated APOE RNA levels in a SORL1-dependent manner. These studies provide a mechanistic link between two of the strongest genetic risk factors for AD.

## Introduction

Alzheimer’s disease (AD) is a progressive neurodegenerative disease that is characterized by extracellular amyloid-beta (Aβ) plaques and intracellular tau tangles. Early onset AD (EOAD) accounts for less than 5% of AD, while late onset AD (LOAD) accounts for the majority of the disease cases. In EOAD, mutations in amyloid precursor protein (APP) or presenilin 1 and 2 (PSEN1 and PSEN2) impair APP processing, which results in accumulation of Aβ. Recent studies have identified rare loss-of-function mutations in SORL1 that appear to be causative for AD, establishing SORL1 as a top AD gene (Pottier *et al*., 2012, Vardarajan *et al*., 2015, Holstege *et al*., 2017, Raghavan *et al*., 2018, Verheijen *et al*., 2016, Thonberg *et al*., 2017, Gomez-Tortosa *et al*., 2018, Le Guennec *et al*., 2018).

Large-scale analyses of genome wide association studies (GWAS) have identified more than 30 loci associated with AD risk (Kunkle *et al*., 2019, Wightman *et al*., 2021, Bellenguez *et al*. 2022). The strongest genetic risk factor for LOAD is the ε4 haplotype of apolipoprotein E (APOE), a protein that has a primary function in cholesterol metabolism (Strittmatter *et al*., 1993, Holtzman, Herz and Bu, 2012, Butt *et al*., 2022). In addition to its role in cholesterol transport, APOE has been implicated in a variety of AD-relevant processes including oligomerization of Aβ, enhancement of neurodegeneration in response to pathologic tau, impairment in endocytic trafficking, and dysfunction of microglia (Shi *et al*., 2017, Wang *et al*., 2021, Chen *et al*., 2021, Chen *et al*., 2010 Xian *et al*., 2018, reviewed in Martens *et al*., 2022 and Lane-Donovan and Herz, 2017). Recent studies have implicated a neuron-specific role for APOE in immune response pathways (Wang *et al*., 2018, Zalocusky *et al*., 2021). Each of these processes likely contributes to AD pathogenesis, but there are still many questions to be addressed regarding APOE biology in the brain. Interestingly, variants in clusterin (CLU, also called apolipoprotein J), another apolipoprotein, also are associated with AD risk.

SORL1 is a type-1 transmembrane receptor that belongs to both the vacuolar protein sorting 10 (VPS10p) containing receptor family and the low-density lipoprotein (LDL) receptor family. Single nucleotide polymorphisms (SNPs) at the SORL1 locus are associated with AD in both candidate gene-based and genome wide association studies (GWAS) (Rogaeva *et al*., 2007, Bettens *et al*., 2008, Miyashita *et al*., 2013, Lambert *et al*., 2013, Kunkle *et al*., 2019, Wightman *et al*., 2021, Bellenguez *et al*., 2022). Importantly, several SORL1 coding variants have been identified in a subset of EOAD patients, (Pottier *et al*., 2012, Vardarajan *et al*., 2015, Holstege *et al*., 2017, Raghavan *et al*., 2018) and variants leading to premature termination codons are found exclusively in AD cases (Holstege *et al*., 2017). In addition, variants that result in truncation of SORL1 have been shown to reduce SORL1 levels in AD individuals (Grear *et al*., 2009, Campion, Charbonnier and Nicolas, 2019). Together, these data suggest that SORL1 is a causal gene for AD (Scheltens *et al*., 2021). Further, reduced expression of SORL1 has been observed in the CSF and post-mortem brain of individuals with late-onset Alzheimer’s disease (Scherzer *et al*., 2004, Dodson *et al*., 2006, Ma *et al*., 2009).

SORL1 has a major role in intracellular trafficking of various cargos, including APP, which is mediated in part through its role in retromer complex-mediated trafficking (Small and Gandy, 2006, Rogaeva *et al*., 2007, Nielsen *et al*., 2007, Fjorback and Andersen, 2012a, Dumanis *et al*., 2015). In the absence of SORL1, APP fails to be transported out of the endosome to the trans Golgi network, cell surface, and lysosomes. This retention of APP in endosomes leads to an increase in generation of Aβ (Andersen *et al*., 2005, Offe *et al*., 2006, Young *et al*., 2015, Andersen, Rudolph and Willnow, 2016). SORL1 also has been shown to directly bind to Aβ and affect its lysosomal degradation (Caglayan *et al*., 2014). In iPSC-derived neurons, loss of SORL1 leads to impaired endosomal trafficking and autophagy, and ultimately increased Aβ levels (Knupp *et al*., 2020, Hung *et al*., 2021). Consistent with these human findings, in AD mouse models, loss of SORL1 leads to an increase in Aβ levels in the brain (Dodson *et al*., 2008).

SORL1 is also referred to as LR11, or LDL receptor with 11 class A ligand binding repeats, and the presence of these ligand binding repeats suggests it also may function as a lipoprotein receptor. Previous studies have shown that SORL1 can regulate the uptake of various lipoproteins in CHO cells (Taira *et al*., 2001), and exogenously overexpressed SORL1 in HEK293 cells can bind to APOE in an isoform-dependent manner (Yajima *et al*., 2015). However, to understand if and how SORL1 and APOE may interact to affect AD risk, it is important to use a more physiologically relevant system wherein SORL1 is expressed at endogenous levels in cell types found in the brain. Moreover, the physiological consequences of SORL1 and APOE interaction, if any, are unknown, and it is unclear if such consequences are relevant to the known role of SORL1 in the endolysosomal trafficking pathway and/or to other unidentified functions of SORL1.

Here, we aimed to understand the intersection of SORL1 and APOE biology in human brain cells. Given that SORL1 is highly expressed across a variety of cell types, we performed transcriptomic profiling comparing wild type to SORL1 null iPSC lines that were differentiated to a variety of brain cell types (neurons, astrocytes, microglia, endothelial cells). This screen revealed that neurons and astrocytes are most sensitive to loss of SORL1, and that overlapping and distinct pathways are altered in different cell types. We found that loss of SORL1 leads to a significant reduction of both extracellular and intracellular APOE protein levels specifically in neurons but not in astrocytes or microglia. A similar reduction was observed in neurons in another lipoprotein associated with AD, CLU. In addition, loss of SORL1 in neurons led to an increase in Aβ levels and phosphorylation of tau. Interestingly, while Aβ and p-tau elevation are rescued by enhancing retromer or autophagy function in SORL1 null neurons, reduced APOE levels were not altered, suggesting that APOE regulation is a separable phenotype from APP processing and abnormal tau phosphorylation in SORL1 loss-of-function. We further interrogated the relationship between APOE, CLU and SORL1 in a set of 50 iPSC lines derived from the ROS and MAP aging cohorts. Intriguingly, following differentiation, a strong correlation between naturally varying levels of SORL1 and both APOE and CLU expression was observed in neurons but not in astrocytes. Analyses of snRNAseq directly from brain tissue from ROS and MAP participants again revealed a strong association of SORL1 with both APOE and CLU expression within neurons, supporting the relevance of this relationship in the aged human brain. Pathway analyses of multiple datasets implicated TGF-β/SMAD signaling in SORL1 biology, and stimulation and inhibition of this pathway modulated APOE RNA and protein levels. By incorporating multiple iPSC experimental systems together with post-mortem brain snRNAseq data, we demonstrate a potential mechanism linking two strong AD risk genes, SORL1 and APOE, that occurs selectively in neurons.

## Results

### Astrocytes and neurons are most sensitive to loss of SORL1

SORL1 is expressed in many cell types of the brain, and an unanswered question in the field is which cell type(s) are most sensitive to a reduction in levels of SORL1. Here, we performed an RNA sequencing (RNAseq) screen to compare gene expression in SORL1 knock out (KO) cells to wild type (WT) cells across cell types. We utilized a previously described pair of wild type and SORL1 homozygous KO iPSC lines derived from a healthy male (Knupp *et al*., 2020). Paired WT and KO iPSCs were differentiated into neurons (iNs), astrocytes (iAs), microglia (iMGLs), and endothelial-like cell fates (iBMVECs) following previously published protocols (Fig. 1A, Supp. Fig. 1A-D). In brief, pure (>95%) cultures of iNs or iAs were generated using transduction of lentivirus expressing either NGN2 (a transcription factor that induces neuronal fate), or both SOX9 and NFIB (transcription factors shown to induce astrocyte fate), using a Tet-inducible system (Zhang *et al*., 2013, Canals *et al*., 2018). These protocols generate brain cell types that have been extensively characterized previously (Muratore *et al*., 2017, Lagomarsino *et al*., 2021, Gao *et al*., 2019, Dobrindt *et al*., 2020). iMGLs were differentiated using a protocol that utilizes a hematopoietic stem cell intermediary to generate microglia-like cells, >95% of which express the microglial marker IBA1, as previously described (Abud *et al*., 2017, McQuade *et al*., 2018). iBMVECs were generated by initial co-differentiation into neural and endothelial progenitors and then subsequent selection and differentiation into endothelial-like cells (Lippmann *et al*., 2012, Lippmann *et al*., 2015, Park *et al*., 2019). Once the differentiations were complete (at iN d21, iA d21, iMGL d40, and iBMVEC d14), samples were harvested for transcriptomic analysis and immunostaining. As previously described, cell cultures expressed expected cell-type-specific markers, as assayed by immunostaining and RNAseq (Fig. 1B,C).

**Figure 1.**
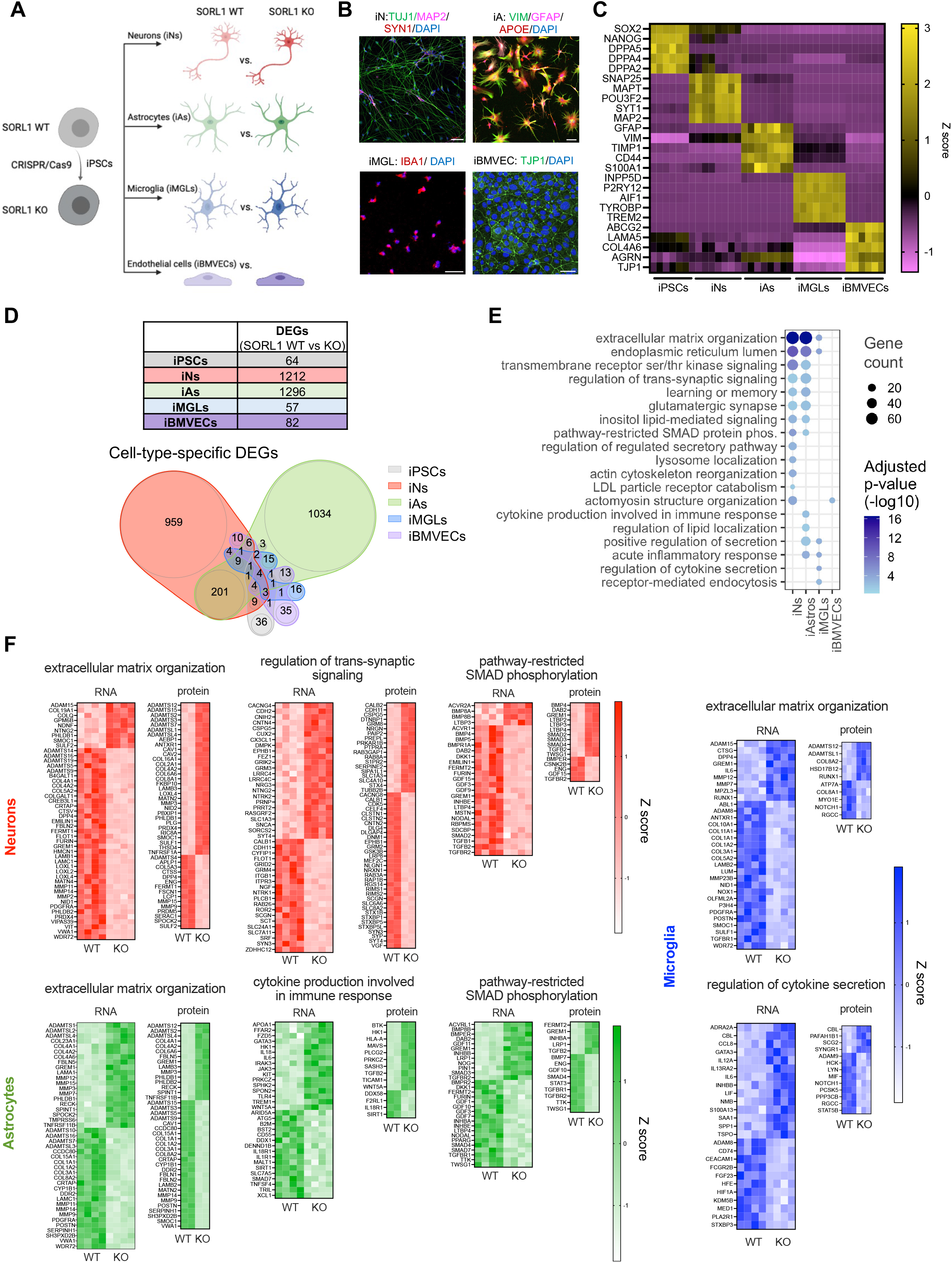
Astrocytes and neurons are most sensitive to loss of SORL1. (A) Schematic of the experimental design. SORL1 KO iPSCs were generated using CRISPR/Cas9 as previously published (Knupp et al. 2020), and paired KO and WT iPSCs were differentiated into neurons (iNs), astrocytes (iAs), microglia (iMGLs), and endothelial cells (iBMVECs). Schematic was made using Biorender. (B) Representative immunocytochemistry images of iN, iA, iMGL, and iBMVEC cultures. Scale bars = 50um. (C) Heatmap of RNA expression (Z score) of cell-type-specific markers. (D) Cultures were analyzed via RNAseq, and differential expression analyses performed within each cell type. DEGs were identified using a Wald test (FDR q-value < 0.05 and b > 0.5). The number of differentially expressed genes between SORL1 WT and KO in each cell type are shown. The five-way venn diagram demonstrating overlap of common DEGs between cell types was generated using nvenn (Pérez-Silva, Araujo-Voces and Quesada, 2018). All significant pathways had a q-value <0.05. (E) Dot plot showing significant overrepresented gene ontology (GO) pathways in iNs, iAs, iMGLs and iBMVECs. Dots are sized based on the number of pathway factors in DEGs and colored by adjusted p-value. (F) Proteomic profiling using TMT-MS was performed in iN, iA, and iMGL cultures. Shown are heatmaps of differentially expressed genes (DEGs) and differentially expressed proteins (DEPs) for a subset of pathways enriched in iNs, iAs, and/or iMGLs. See Supp. Tables 1 and 2 for full data set.

To determine which cell type is most sensitive to loss of SORL1, we performed differential expression analyses to compare WT to SORL1 KO cells of each fate. We used cutoffs of q-values < 0.05 and absolute value of b (fold change) > 0.5 to identify differentially expressed genes (DEGs). Neurons and astrocytes showed the highest number of DEGs (1212 and 1296 respectively) compared to iPSCs, microglia, and endothelial cells (Fig. 1D, Supp. Table 1). Enrichment analyses identified pathways that are differentially enriched in each cell type (Supp. Table 2). Some pathway alterations were shared across cell types while others were specific to a single cell type (Fig. 1E). In parallel to RNAseq analyses, unbiased proteomic profiling also was performed in each cell type to identify individual factors and pathways that had similar responses to SORL1 knockout at both the RNA and protein level. Pathways consistently altered in both RNA and protein data sets include “extracellular matrix organization” between neurons, astrocytes, and microglia, “pathway-restricted SMAD phosphorylation” between neurons and astrocytes, and gene sets relating to cytokine production between astrocytes and microglia (Fig. 1F). Cell-type-specific enrichment was observed in terms such as “lysosome localization” in neurons and “regulation of lipid localization” in astrocytes (Fig. 1E,F). These analyses provide a first view into the shared and cell-type-specific genes and pathways altered with SORL1 loss-of-function across human brain cell types.

### APOE and CLU are reduced in SORL1 KO neurons

In addition to pathway analyses, we investigated whether any LOAD GWAS genes were differentially expressed in the RNAseq data. Intriguingly, several LOAD genes showed disrupted expression in SORL1 null iNs, iAs, or iMGLs (Fig. 2A, Supp. Fig. 2A-J). Due to the introduction of a premature stop codon, nonsense mediated decay results in reduced SORL1 levels across all cell types (Fig. 2B). Microglia expressed the highest RNA levels of SORL1 in WT cells, and consequently the highest levels of RNA in the KO (likely due to the rates of nonsense-mediated decay). Although some RNA was detected in KO cells, protein levels were undetectable by WB in all cell types (Fig. 2G-I). CD2AP and BIN1, both genes with known trafficking roles, and two apolipoproteins, APOE and CLU (APOJ), showed cell-type-specific changes in their expression levels (Fig. 2C-F). With the loss of SORL1, APOE and CLU RNA levels decreased in neurons, but increased in astrocytes (Fig. 2E-F). APOE and CLU were not affected by SORL1 loss in iPSCs, microglia, and endothelial cells, suggesting cell-type-specific relationships. In SORL1 KO neurons, RNA levels of additional apolipoproteins were reduced including APOC1 and APOL4, while in SORL1 KO astrocytes, APOE and CLU were the only apolipoproteins elevated as a result of loss of SORL1 (Supp. Table 1).

**Figure 2.**
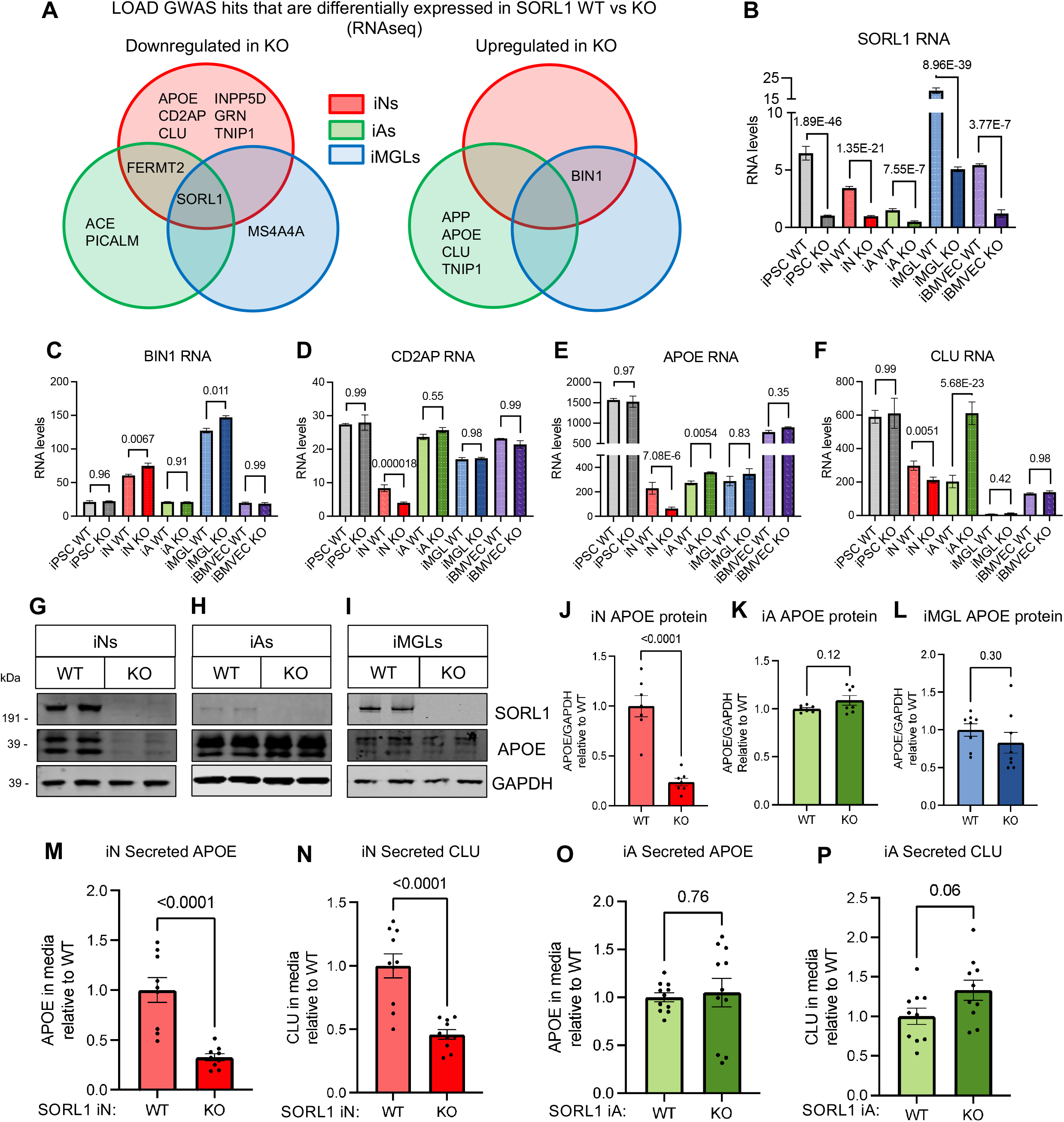
APOE and CLU are reduced in SORL1 KO neurons. (A) Venn diagram of LOAD GWAS genes that are differentially expressed in SORL1 KO neurons, astrocytes, and microglia. Genes with p-adj < 0.05 were included in the diagram. (B-F) SORL1, BIN1, CD2AP, APOE, and CLU RNA levels comparing SORL1 WT versus KO across cell types. (G-I) Representative western blot of iNs, iAs, and iMGLs showing protein expression levels of SORL1, APOE, and GAPDH. (J-L) Quantification of APOE/GAPDH in iNs, iAs, and iMGLs. (M-P) Secreted APOE and CLU values were measured using MSD ELISA, and the values were normalized to WT within each comparison. Data show mean +/- SE from 3 differentiations, n=3-4 per differentiation for each line. Unpaired student’s t test (two tailed).

Consistent with the RNAseq results, APOE protein levels were significantly decreased in SORL1 KO neurons (Fig. 2G,J). In comparison, there were no detectable differences in APOE protein levels in SORL1 KO astrocytes or microglia (Fig. 2H,I,K,L). Since APOE is a secreted protein, it was possible that APOE in the iN cell lysate could be decreased due to an elevation in secretion. However, APOE levels in the media also showed a dramatic reduction (Fig. 2M). The impact of loss of SORL1 was not isolated to APOE, as secreted CLU levels also were reduced in SORL1 KO iNs (Fig. 2N). Secreted APOE and secreted CLU were not significantly changed in SORL1 KO astrocytes, indicating that this SORL1 regulation of APOE and CLU is neuron-specific (Fig. 2O,P). This cell-type specificity of regulation could explain why changes in APOE and CLU have not been previously reported, and underscores the importance of studying the cell-type-specific roles of AD genes of interest.

### SORL1 KO neurons show elevated Aβ and increased phosphorylation of tau

To test the reproducibility of finding of SORL1 KO in d21 iNs, we used CRISPR/Cas9 editing to generate a second isogenic pair of SORL1 null iPSCs (Fig. 3A). The second cell line was generated from a previously described female iPSC line (Muratore *et al*., 2017). A gRNA targeting a different exon was employed to generate cell lines with an indel introduced at the SORL1 locus that results in nonsense mediated decay (Supp. Fig. 3A-C). The two pairs of WT and SORL1 KO iPSC lines were differentiated into neurons (iNs) following the NGN2 protocol, and all four lines showed robust differentiation to neuronal fate with no major differences in morphology as depicted by immunostaining images with the neuronal markers TUJ1, POU3F2, and MAP2 (Fig. 3B). iN lysates from both cell line pairs 1 and 2 show that protein expression of SORL1 is eliminated in SORL1 KO lines (Fig. 3C). Consistent with our observations in line 1 iNs, APOE protein levels were dramatically reduced in SORL1 KO iNs in line 2 (Fig. 3D). Secreted APOE and CLU levels also were reduced in SORL1 KO iNs from cell line 2 (Fig. 3E,F). These data in a second genetic background strengthen the findings that SORL1 regulates levels of neuronal APOE and CLU.

**Figure 3.**
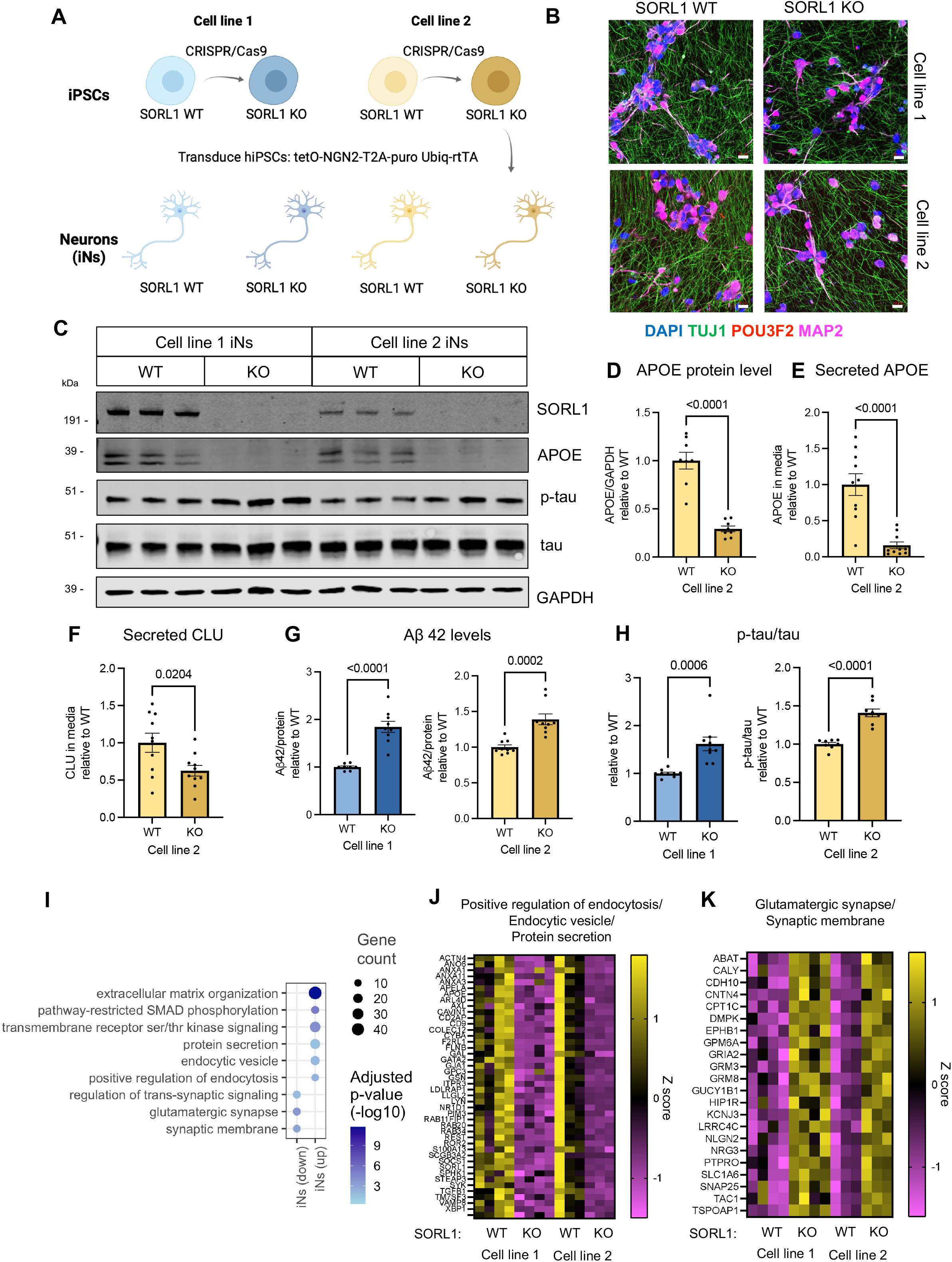
SORL1 KO neurons express elevated Aβ and increased phosphorylation of tau. (A) Overview of the experimental design to generate SORL1 KO neurons. SORL1 KO iPSCs were generated using CRISPR/Cas9 and were differentiated into neurons using induced-expression of NGN2. Schematic generated using Biorender. (B) Representative immunocytochemistry images of d21 iNs showing the expression of neuronal markers. Scale bars = 20um. (C) Representative western blot of iN d21 protein lysates from line 1 and line 2. Quantification of APOE/GAPDH in lysates (D), APOE in the media (E) and CLU in the media (F) normalized to total protein for cell line 2 iNs. (G) Aβ42 levels in the media normalized to protein concentration. (H) Quantification of p-tau (p202/205)/total tau ratio in iN d21 protein lysates from western blot data. SORL1 KO iNs from line 1 and 2 both show increased phosphorylation of tau. Data in D-H show mean +/- SEM from three differentiations, n=3 per differentiation for each line. Values are normalized to WT. Unpaired student’s t test (two tailed). (I) Dot plot of molecular pathway over-representation in up or down-regulated DEGs in both lines 1 and 2. All pathway enrichments have an adjusted pval (q) <0.05). See also Supp Table 3 for full dataset. (J-K) Heatmap of a subset of genes that are differentially expressed in SORL1 WT vs KO iNs and their relative expression levels (Z-score) from selected GO terms in (I).

Accumulation of Aβ and tau are the two major histopathological hallmarks of AD. Multiple studies have shown that APP is one of the cargos of SORL1 in its neuronal retromer role and loss of SORL1 leads to retention of APP in endosomes, resulting in an increase in Aβ generation (Offe *et al*., 2006, Dodson *et al*., 2008, Knupp *et al*., 2020). As predicted, SORL1 KO iNs show an increase in Aβ42 generation (Fig. 3G). All three Aβ peptides detected by MSD ELISA (38, 40, and 42) were increased in SORL1 KO iNs, without alteration of the Aβ42/40 ratio or detectable protein expression of full-length APP, suggesting that SORL1 affects the generation of Aβ but does not alter the processivity of gamma-secretase (Supp. Fig. 4A-I).

**Figure 4.**
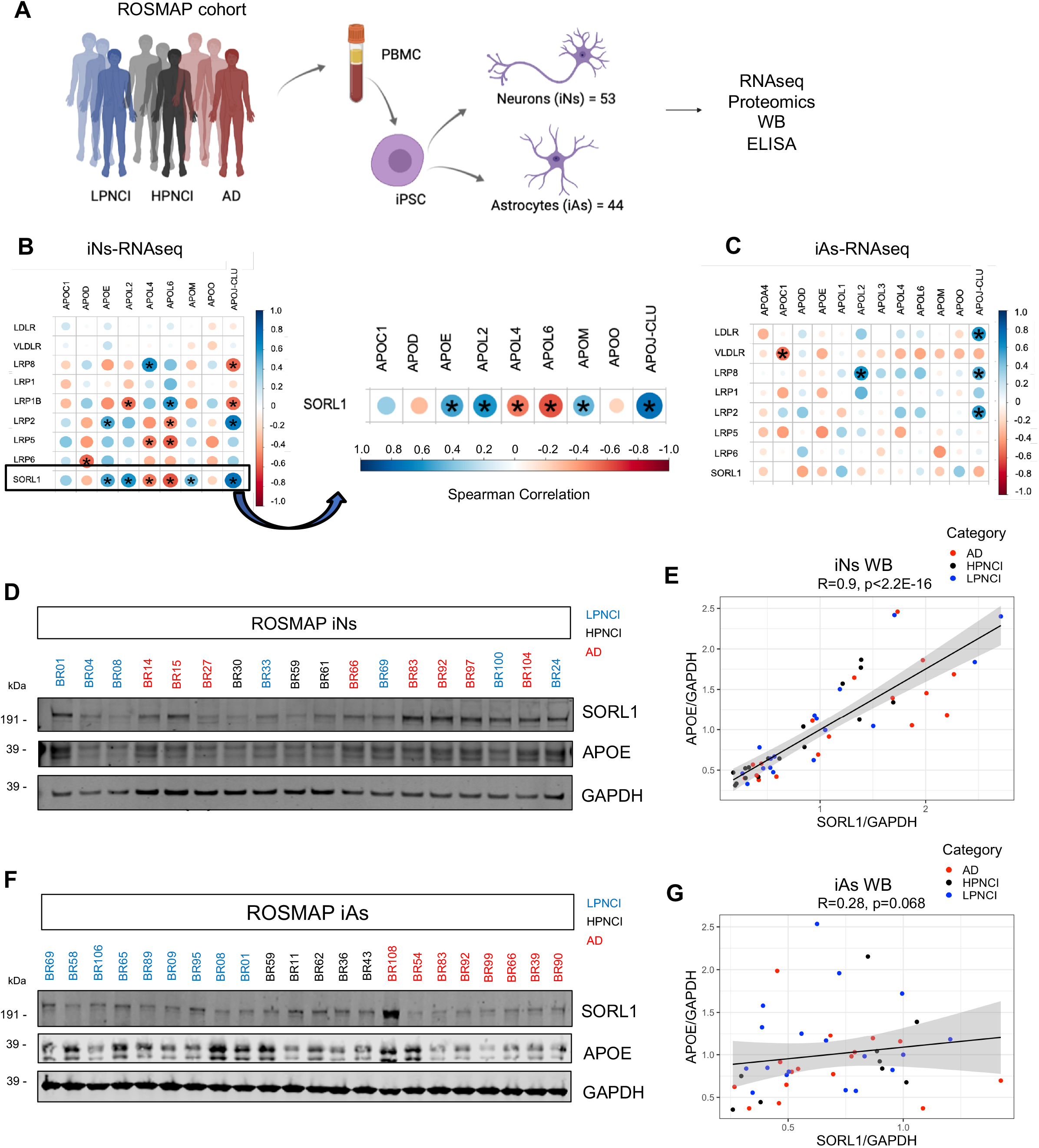
SORL1 shows a strong positive correlation with APOE and CLU in iPSC-derived neurons from the ROSMAP aging cohort. (A) PBMCs were collected from ROSMAP cohort participants, which were converted to iPSCs and differentiated into neurons (iNs) and astrocytes (iAs). Schematic was generated using Biorender. (B-C) The expression levels of SORL1 and other members of the LDL receptor family were each compared to APOE and members of the apolipoprotein family in both iNs (B) and iAs (C). Correlation dot plots are used to represent Spearman correlations for each comparison with the size of the dot increasing with increasing with lower p values (asterisks indicate pval < 0.001) and color indicating r value (blue for positive and red for negative r values). (D) A representative western blot image of SORL1, APOE, and GAPDH expression in ROSMAP iN protein lysates. Individuals have a unique ID that is colored based on their AD diagnosis status. LPNCI: low AD neuropathology not cognitively impaired, HPNCI: high AD neuropathology not cognitively impaired and AD: Alzheimer’s disease. (E) Spearman correlation between SORL1/GAPDH and APOE/GAPDH levels in iN lysates. r = 0.90, p = 2.2e-16. (F) A representative western blot image of SORL1, APOE, and GAPDH expression in ROSMAP iA protein lysates. (G) Spearman correlation between SORL1/GAPDH and APOE/GAPDH levels in iA lysates. r = 0.28, p = 0.068.

The second neuropathological hallmark of AD is the accumulation of intracellular tau tangles. Prior to overt tangle formation, an elevation of phosphorylation of tau occurs. SORL1 KO iNs showed an increase in phosphorylated tau levels (as measured by AT8, which recognizes phosphorylated serine residues at positions 202 and/or 205) relative to total tau (Fig. 3H). To our knowledge, this is the first report of modulation of phospho-tau levels in SORL1 null human neurons.

We next investigated whether APOE genetic status affects SORL1 levels. We examined neurons derived from an isogenic set of APOE KO, APOE ε4/ε4, and ε3/ε3 iPSCs. SORL1 protein levels and p-tau/tau levels were not altered across these iNs (Supp. Fig. 5A-C). Additionally, we used two different APOE shRNA constructs to knock down APOE expression in INs. These constructs successfully reduced the APOE levels to ∼30%, however, SORL1 levels were not altered (Supp. Fig. 5D-G). Importantly, p-tau levels were not altered in APOE KO neurons, suggesting that the reduction in APOE observed in SORL1 KO iN is not sufficient to induce the p-tau phenotype.

**Figure 5.**
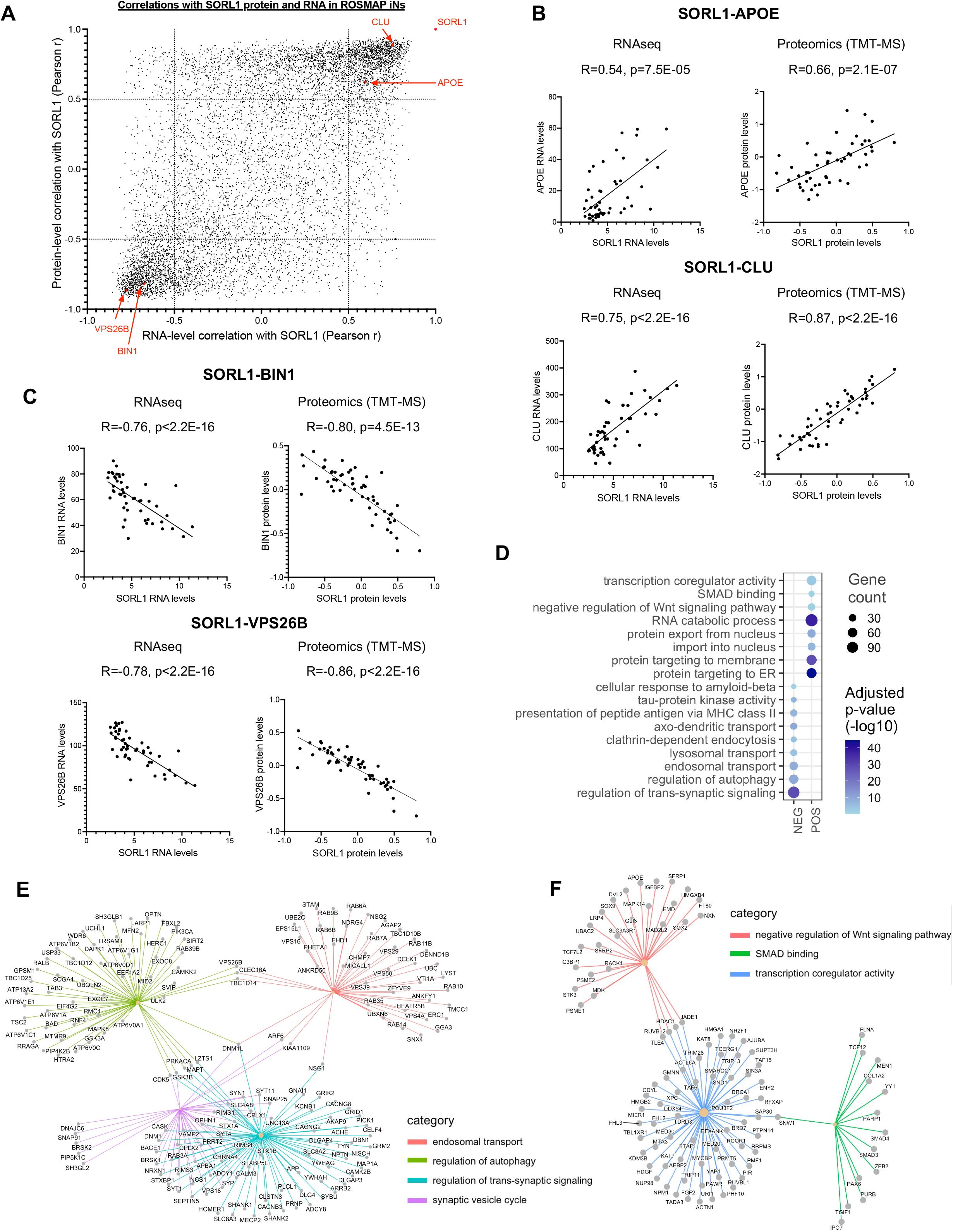
Unbiased RNAseq and proteomics of ROSMAP iNs identify genes and pathways associated with genetically encoded natural variation in SORL1 levels. **(A)** ROSMAP iPSC lines were differentiated to neuron fates in independent differentiations, and RNAseq and TMT-MS proteomic profiling performed. Pearson correlation coefficients between SORL1 and all other factors detected within both the RNAseq and proteomics data sets are plotted. Each dot represents data for a single gene. High density areas of the plot illustrate factors that are either positively (upper right section) or negatively (lower left section) correlated with SORL1 in both the RNAseq and the proteomics data. Points representing selected genes of interest (SORL1, APOE, CLU, BIN1, and VPS26B) are highlighted with red arrows. (B) Correlation between SORL1 and APOE, CLU in ROSMAP iN RNAseq and proteomics data. Each dot represents data from iNs derived from one individual. (C) Correlation between SORL1 and BIN1 and VPS26B in ROSMAP iN RNAseq and proteomics data. (D) Pathway enrichment dot plot representing biological gene cohorts that are significantly overrepresented in either the upper right (POS) or lower left (NEG) sections of the SORL1 correlation plot in (A). Dot size represents the number of genes passing correlation cutoff that were in the gene set while color indicates q-value. All pathways shown had a q-value for enrichment of 0.02 or better. (E,F) Gene concept network plot representing genes that belong to the pathways identified in (D) that are driving the positive or negative associations with SORL1 expression.

### Loss of SORL1 in human neurons induces a downregulation of genes associated with endocytosis, SMAD signaling, and ECM organization and an upregulation of genes encoding synaptic proteins at the RNA level

To identify pathways affected by loss of SORL1 and therefore potentially contributing to the increase in phosphorylated tau and reduction of APOE in SORL1 KO iNs, genes were identified that were concordantly differentially expressed between SORL1 WT and KO in both genetic backgrounds (q < 0.05 for inclusion) (full data set in Supp. Table 3). Gene ontology analysis revealed biological processes associated with SORL1 levels including terms also identified in the initial screen of DEGs across cell types such as “extracellular matrix organization”, “regulation of pathway restricted SMAD protein phosphorylation”, and “glutamatergic synapse”, as well as “regulation of endocytosis”, “endocytic vesicle”, and “protein secretion” (Fig. 3I-K). Importantly, each of the terms listed in Fig. 3I also were replicated in an independently published RNAseq data set of SORL1 KO vs WT neurons, which was generated using a different, undirected protocol for differentiation to neuronal fate (Mishra *et al*., 2022; Supp. Table 4). SORL1 dependent effects on trafficking genes are consistent with its known role in neurons in retromer-mediated trafficking (Fjorback *et al*., 2012b, Small and Petsko, 2015) and the analyses provide new insights into the specific genes altered with SORL1 loss-of-function. The pathways affected by loss of SORL1 may result from disruption of SORL1’s function in retromer-related endosomal trafficking, its potential role as a lipoprotein receptor, or other unidentified molecular interactions, and the cellular systems described here provide a resource for interrogating these candidate mechanisms. Analyses of complementary datasets in the following sections aid in our selection of robust candidate pathways to interrogate mechanistically in the final section.

### SORL1 shows a strong positive association with APOE and CLU in iPSC-derived neurons from the ROSMAP aging cohorts

SORL1 levels have been reported to be reduced in the AD brain, and rare truncation variants have been identified in the heterozygous state that are predicted to be loss-of-function (Scherzer *et al*., 2004, Grear *et al*., 2009, Pottier *et al*., 2012, Vardarajan *et al*., 2015, Cuccaro *et al*., 2016, Verheijen *et al*., 2016, Campion, Charbonnier and Nicolas, 2019). While the KO vs WT experimental system is valuable for uncovering cellular roles of SORL1, complete loss of SORL1 does not accurately represent a state that occurs in the human brain. In order to probe the relationship between SORL1 and APOE and other apolipoproteins in a system that captures genetically encoded variation in SORL1 levels in human neurons, we utilized a cohort of >50 iPSC lines derived from the ROS and MAP cohorts (Lagomarsino *et al*., 2021). The ROS and MAP (Religious Order Studies and Rush Memory and Aging Project) cohorts are composed of individuals ranging in age from nearly 60 to over 100 years old who span the range of neuropathological and clinical outcomes (Bennett *et al*., 2018). Previously, we reported the generation of these iPSC lines and showed that iNs from these individuals reflect significant features of the protein networks and neuropathology present in the brain tissue of the same individuals from whom the cells were derived (Lagomarsino *et al*., 2021). Here, we utilized this system to interrogate putative connections between apolipoproteins and SORL1. First, ∼50 individuals (53 for iNs, 44 for iAs) were selected to reflect varying degrees of pathological and clinical diagnosis, then iPSCs from selected individuals were differentiated into neuron (iN) and astrocyte (iA) fates following the same differentiation protocols described previously (Fig. 4A; Supp. Fig. 6A-F).

**Figure 6.**
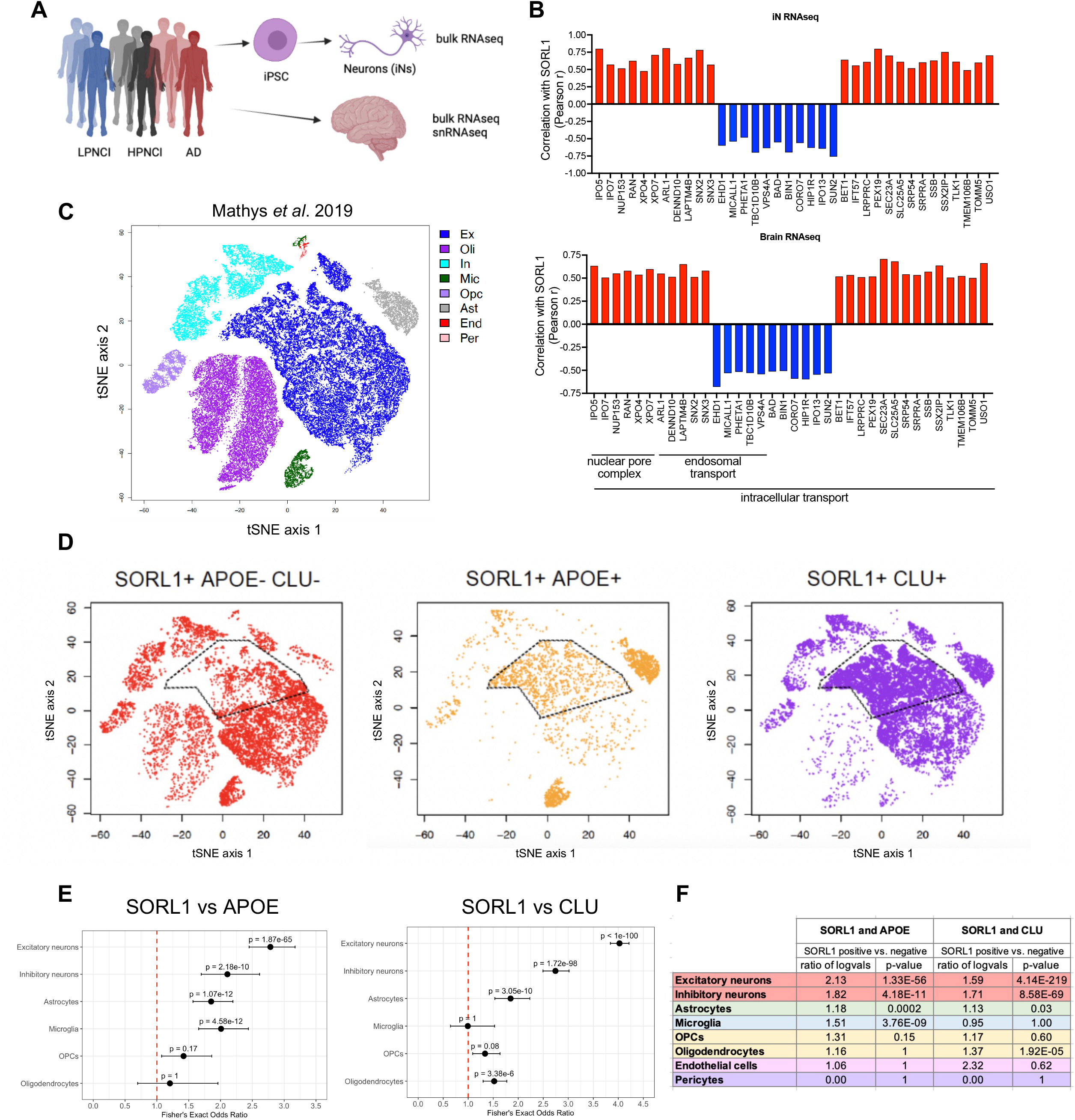
snRNAseq of post-mortem brain tissue validates an association between SORL1 and both APOE and CLU in excitatory neurons. (A) iN bulk RNAseq and post-mortem brain (medial frontal cortex) bulk RNAseq as well as snRNAseq dataset were generated from the ROSMAP cohort. Schematic was generated using Biorender. (B) Correlation coefficient values (Pearson r) were calculated between SORL1 and other genes in bulk RNAseq data from iNs and brain tissue. Concordant associations of genes in intracellular transport, endosomal transport, and nuclear pore complex gene sets within iN and brain RNAseq datasets. (C) TSNE plots of snRNAseq data derived from the ROSMAP cohort, colored by cell type, as determined by marker expression (Mathys et al. 2019). Data are from 70,634 cells from 48 individuals. (D) TSNE plots with cells colored by SORL1, APOE and/or CLU detection, as labelled. High overlap between SORL1 expression and APOE and CLU expression in a subset of excitatory neurons. (E) Fisher’s exact test results (adjusted p-value and odds ratio) for the detection overlap of SORL1, APOE, and CLU within nuclei, separated by cell type. (F) Differential expression results of APOE and CLU between SORL1+ and SORL1-nuclei using a Wilcoxcon rank sum test, separated by cell type. See also Supp. Table 5.

Analysis of RNAseq data across these ROSMAP iNs and iAs revealed a high correlation between APOE and CLU with SORL1 in neurons but not astrocytes (Fig. 4B,C). CLU and APOE also showed robust positive correlations with SORL1 in iNs at the protein level, as measured by western blot (APOE) (Fig. 4D,E) and ELISA of the conditioned media (APOE, CLU) (data not shown). As in the RNAseq data, western blot and ELISA analyses of iAs derived from ROSMAP iPSCs showed no such correlation between APOE or CLU and SORL1 (Fig. 4F,G). The iN experimental system captures the genetics of the individuals from whom the cells are derived, and varying levels of SORL1 and APOE are encoded at least in part within the genome of the individuals. It is important to note that while SORL1 5’ haplotypes (Rogaeva *et al*., 2007, Young *et al*., 2015) and APOE haplotype contribute to the RNA and protein levels across iNs, they do not fully explain the variance in levels of SORL1 and APOE (Supp. Fig. 7A-H). These data suggest that other genetic variants are contributing in a complex manner to affect levels of SORL1, which in turn affects APOE levels, and future studies with greater power will be required to uncover the genetic variants that are driving varying levels of SORL1.

**Figure 7.**
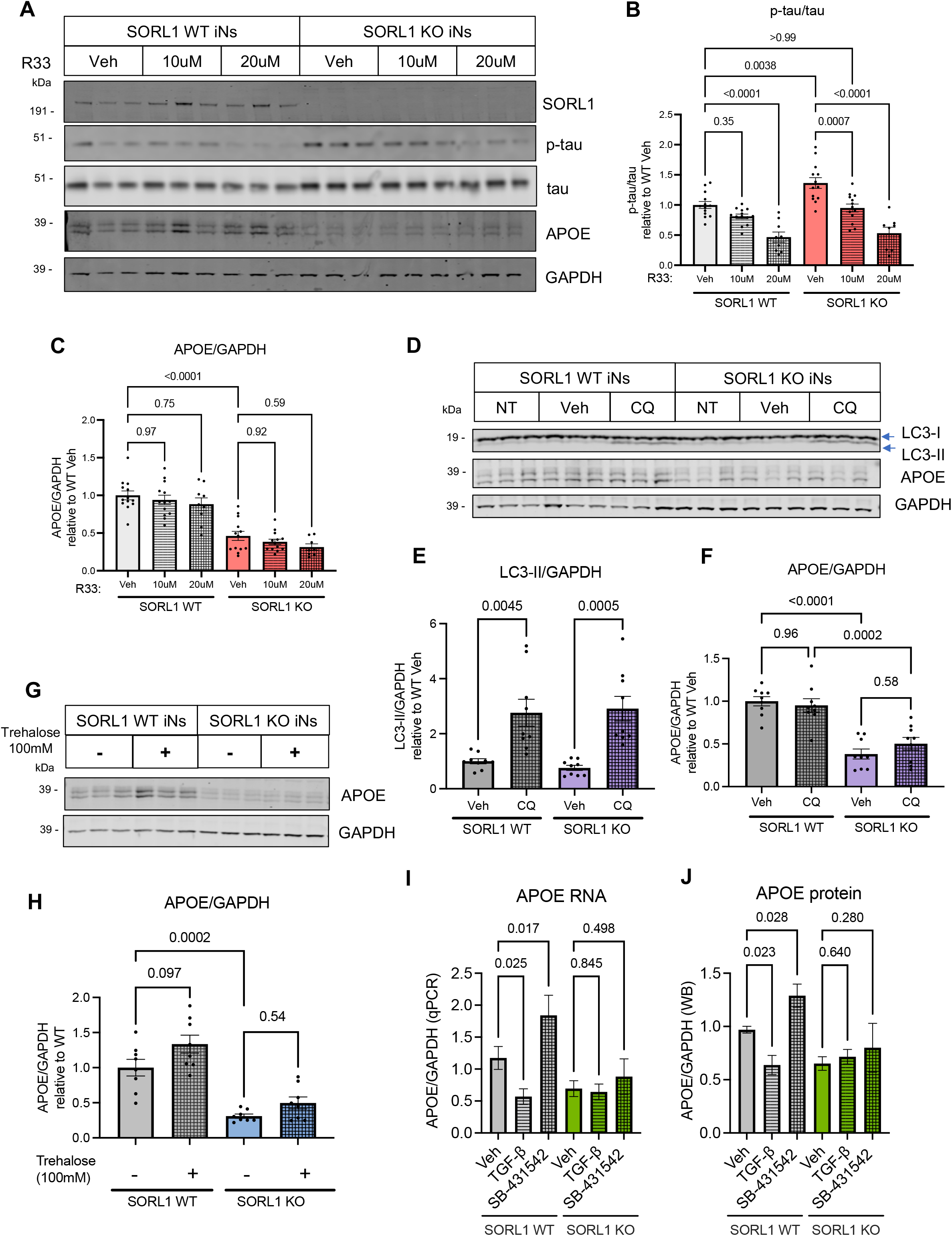
Enhancement of retromer function or autophagy rescues Aβ and tau phenotypes while modulation of SMAD signaling regulates APOE levels in a SORL1 dependent manner. (A) A representative western blot of SORL1 WT and KO iNs after R33 treatment. INs were treated with R33, a retromer stabilizer, at either 10uM or 20uM for 72hrs, and harvested at D21 of differentiation. Quantification of the protein expression of (B) p-tau/tau (p202/205), and (C) APOE/GAPDH. (D) A representative western blot of APOE, GAPDH, and LC3-I and LC3-II after 72hrs of 5uM chloroquine (CQ) treatment. (E-F) Quantification of LC3-II/GAPDH and APOE/GAPDH. (G) A representative western blot of APOE and GAPDH after trehalose treatment. iNs were treated with trehalose, a putative enhancer of autophagy, at 100mM for 72hrs and harvested at d21 of differentiation. (H) Quantification of the protein expression of APOE/GAPDH. Data show mean +/- SEM from three differentiations, n=3 per differentiation for each line. Values are normalized to WT iNs treated with vehicle. One-way ANOVA with Tukey’s multiple comparisons test. (I,J) SORL1 WT and KO iNs (line #1) were treated with vehicle, TGF-β or SB-431542 for 72 hours. RNA was collected and qPCR performed for APOE and GAPDH (I) or else cells lysed and western blotting performed for APOE and GAPDH (J). For (I,J) data show mean +/- SEM from three differentiations, n=2-4 wells per differentiation for each line. Values are normalized to WT iNs treated with vehicle within each experiment. One-way ANOVA with Fisher’s exact test.

In addition to APOE and CLU, SORL1 levels were significantly positively correlated with a subset of other apolipoproteins in neurons (APOL2 and APOM). However, the positive associations of SORL1 with apolipoproteins was not universal: SORL1 levels were *negatively* correlated with APOL4 and APOL6 levels (Fig. 4B). In addition, the association of lipoprotein receptors with APOE and CLU was specific to SORL1 and LRP2 in neurons, with other related lipoprotein receptors showing no such association (Fig. 4B).

### Unbiased RNAseq and proteomics of ROSMAP iNs identify genes and pathways associated with genetically encoded natural variation in SORL1 levels

To globally examine the individual genes and pathways associated with SORL1 in ROSMAP iNs in an unbiased manner, we calculated the correlation coefficients between SORL1 and all factors detected within both the RNAseq and proteomics (TMT-MS) data sets, and plotted protein correlations against transcript correlations for each factor (Fig. 5A). This analysis allowed us to identify factors that were consistently associated with SORL1 levels at both the RNA and protein levels across independent differentiations. While RNA and protein expression do not always agree given the important contributions of post-transcriptional and post-translational regulation, there was clear overall concordance between RNA and protein associations with SORL1 for a subset of genes (Fig. 5A). First, we confirmed that SORL1 shows a strong positive correlation with APOE and CLU in both RNA and protein datasets (Fig. 5B). Several genes of interest to SORL1 biology were identified in this association analysis including the LOAD GWAS gene BIN1 and the retromer protein VPS26B, and these genes showed a negative correlation with SORL1 (Fig. 5C). In accord with observations in SORL1 KO iNs, pathway analyses revealed significant associations between SORL1 levels and intracellular transport mechanisms. For example, SORL1 positively correlated factors were significantly (FDR<0.01) enriched for gene sets involved in protein targeting to the membrane, nucleus, and ER, while SORL1 negatively correlated factors were enriched in gene sets involved in endosomal and lysosomal transport, among others (Fig. 5D). Elevated levels of “tau protein kinase activity” and “cellular response to amyloid beta” also were identified with lower SORL1 levels (Fig. 5D), consistent with the observed increase in Aβ and p-tau observed in SORL1 KO iNs (Fig. 3). Gene concept network (GCN) analyses revealed that SORL1-associated endosomal transport and autophagy pathways are linked to one another via VPS26B, CLEC16A, and TBC1D14, and to synaptic pathways via DNM1L and other genes (Fig. 5E). Similarly, SORL1-associated WNT and SMAD signaling were linked to one another via transcription coregulator activity (Fig. 5F). These data sets and analyses provide a framework in which to further probe the cellular consequences of loss of neuronal SORL1.

### snRNAseq of post-mortem brain tissue validates an association between SORL1 and both APOE and CLU in excitatory neurons

The data described herein suggest that there is a strong relationship between SORL1 and both APOE and CLU that is neuron-specific. To assess if this relationship also exists in the aged human brain, we examined RNAseq data from the medial frontal cortex from the same ROSMAP individuals for whom we derived iPSC lines (Fig. 6A). Not surprising given the cell-type specificity in the cell model, there was no association between SORL1 and APOE at bulk-level in brain tissue. Pathway analysis of genes associated with SORL1 in both the brain data and the iN data (r-value > 0.5) highlighted a set of genes involved in intracellular transport, endosomal transport and the nuclear pore complex that were concordantly associated with SORL1 expression in the brain and iNs (Fig. 6B). Intriguingly, two of these genes, BIN1 and TMEM106B, are LOAD GWAS genes (Fig. 6B).

Finally, to examine cell-type-specific relationships between APOE and SORL1, we examined previously reported single nucleus RNAseq data derived from the ROSMAP cohort (Mathys *et al*., 2019). This dataset is generated from 48 individuals with a median age of 86.9, 24 of which had a diagnosis of AD. 70,634 cells in total were used for the analysis (Fig. 6C). Visual inspection of tSNE plots of the data showed SORL1 expression across virtually all cell clusters. Within the excitatory neuron clusters (Fig. 6C, darkblue), a large proportion appeared to be co-expressing SORL1, APOE, and CLU (Figure 6D). To quantify this result, we performed two types of analysis. First, we performed a Fisher’s exact test to determine whether the percentage of cells expressing SORL1 that express APOE or CLU in each cell type cluster is greater or less than expected by chance (Fig. 6E, Supp. Table 5). For both APOE and CLU, excitatory neurons demonstrated the highest odds ratio, indicating that this cell type is the most enriched for double positive (SORL1+/APOE+ or SORL1+/CLU+) cells. Next, we compared the ratio of log-transformed means for APOE or CLU expression in SORL1 positive versus negative cells. Differential expression analyses show that APOE expression is higher in SORL1+ neurons (2.1-fold for excitatory neurons) (Fig. 6F). Similarly, CLU expression is higher in SORL1+ neurons (1.6-fold for excitatory neurons) (Fig. 6F, Supp. Table 5). Taken together, both methods revealed that expression of SORL1 is strongly associated with both APOE and CLU in excitatory neurons in the human brain.

### Enhancing retromer function via R33 treatment rescues elevated p-tau, but not APOE

We next sought to determine the mechanism by which SORL1 regulates APOE levels in iNs. Unbiased analyses of ROSMAP iNs and SORL1 KO iNs revealed pathways potentially contributing to APOE levels. We first examined whether rescue of defects in retromer-mediated endosomal transport could rescue APOE levels in SORL1 KO iNs. The retromer complex is a multiprotein assembly that is responsible for trafficking cargo from the endosome to the trans-Golgi network and recycling back to the plasma membrane. Recently, it has been shown that in iPSC-derived neurons SORL1 regulates retromer-dependent endosomal recycling in iPSC-derived neurons (Mishra *et al*., 2022). In mice lacking VPS26B, one of the key components of the retromer complex, SORL1 protein levels in the brain were reduced, again supporting a close link between SORL1 and retromer complex (Simoes *et al*., 2021). Analyses of SORL1 loss and natural variation in this study using human cells further supported these findings. To address whether the retromer complex is involved in driving the elevated p-tau or reduced APOE, iNs were treated with R33, a compound that stabilizes the retromer complex by putatively acting as a pharmacological chaperone (Mecozzi *et al*., 2014). It previously has been shown that R33 treatment lowers Aβ generation and phosphorylated tau levels in sporadic AD neurons (Young *et al*., 2018). Here, treatment with R33 at 10uM for 72 hours rescued elevated p-tau/tau in SORL1 KO iNs without altering the p-tau/tau levels of WT iNs. At a higher concentration (20uM), R33 lowered p-tau in both WT and KO iNs (Fig. 7A,B). Furthermore, 20uM R33 treatment partially rescued Aβ levels in SORL1 KO iNs with no change in Aβ42/40 (Supp. Fig. 8A-B) indicating that Aβ and p-tau phenotypes in SORL1 KO iNs are connected to the role SORL1 plays in regulating the retromer complex. However, the same R33 treatments were not sufficient to rescue the decreased APOE levels in SORL1 KO iNs, suggesting that the SORL1 KO APOE phenotype is separable from its Aβ and p-tau phenotypes (Fig. 7C).

### Modulating autophagy via chloroquine or trehalose does not rescue APOE levels

Next, we sought to determine whether the decrease in APOE is a consequence of dysregulated degradation of APOE. Autophagy was associated with SORL1 in the analyses performed herein, and a recent study reported that SORL1 KO neurons show a defect in autophagic flux (Hung *et al*., 2021), raising the possibility that dysregulated autophagy is mediating the effects observed on APOE levels. Alternatively, APOE levels may be regulated by loss of SORL1 via an increase in trafficking of APOE towards lysosomes and the degradation pathway. iNs were treated with 5uM chloroquine or 100mM trehalose at D18 and harvested at D21 for analyses of conditioned media and protein lysates. While chloroquine (CQ) treatment successfully blocked autophagy as measured by an increase in LC3-II protein level (Fig. 7D,E), it did not significantly alter APOE levels in SORL1 WT or KO iNs (Fig. 7F). Next, we treated iNs with trehalose, a putative augmenter of autophagy. After 72hrs of treatment, trehalose reduced the elevated p-tau levels, but did not affect levels of Aβ or APOE in SORL1 WT or KO iNs (Fig. 7G,H, Supp. Fig. 8C-F). Thus, modulation of autophagy by SORL1 loss does not appear to strongly contribute to APOE protein levels in iNs.

### Modulation of SMAD signaling regulates APOE levels in neurons in a SORL1 dependent manner

TGF-β and SMAD signaling pathways were associated with SORL1 through multiple analyses (Fig. 1E,F. Fig. 3I, Fig. 5D,F), and were replicated in an independently generated RNA-seq data set of SORL1 KO iPSC-derived neurons, which utilized an alternative neuronal differentiation protocol (Mishra *et al*., 2022; Supp. Table 4). This association was of particular interest given the observation that SORL1 KO affected APOE RNA levels (Fig. 2E) as well as protein levels (Fig. 2J). SMAD proteins themselves were dysregulated in SORL1 KO iNs (Supp. Fig. 8G-I) and inspection of the APOE genomic locus using the JASPAR database revealed putative SMAD2, SMAD3, and SMAD4 binding sites in the 3 kb region upstream of the APOE transcriptional start site, as well as in the first intron and 3’UTR (Supp. Fig. 8G-I, Supp. Table 6). Analyses of the APOE locus across iNs, iAs, and iMGs via ATAC-seq revealed a region of open chromatin unique to iNs in the promoter, and different degrees of accessibility in additional regions at the APOE locus (Supp. Fig. 8J). To determine if modulation of SMAD signaling affects APOE levels, WT and SORL1 KO iNs were treated either with TGF-β to stimulate SMAD signaling, or else with an inhibitor of TGF-β receptor-mediated SMAD phosphorylation (SB-431542). TGF-β treatment reduced both RNA and protein levels of APOE while SB-431542 increased levels of APOE RNA and protein levels (Fig. 7I,J). This modulation of APOE was SORL1 dependent, as the same treatments had no effects on APOE levels in SORL1 KO iNs (Fig. 7I,J). It is important to note that SMAD signaling is highly complex: SMADs recruit a variety of coactivators and corepressors to genomic DNA to modulate chromatin structure to induce or repress expression at loci of binding (reviewed in Hill, 2016), and future studies will be required to understand the precise mechanism of APOE regulation by TGF-β/SMAD signaling. The results presented here suggest that TGF-β/SMAD signaling acts in concert with other factors to repress APOE transcription in iNs in a SORL1-dependent manner.

## Discussion

SORL1 is expressed across several cell types present in the brain. However, SORL1 functional studies to date have primarily focused on its trafficking role in neurons. Here, we began our study with an unbiased assessment of the consequences of SORL1 loss-of-function across multiple brain cell types. We chose to use the iPSC experimental system for this screen to have a genetically manipulatable human system that can generate an unlimited supply of consistent cell populations. While we found that SORL1 expression was highest in microglia at both the RNA and protein level, the highest numbers of DEGs with SORL1 KO were identified in astrocytes and neurons. While microglia showed many fewer DEGs, this does not preclude an important role for SORL1 in microglia. Indeed, BIN1, a gene consistently associated with LOAD through GWAS, was elevated in SORL1 KO microglia, as were MHC-II genes and other inflammatory response genes (Fig. 1E,F, Fig. 2D, Supp. Table 1). Across cell types, both shared effects of loss of SORL1 and cell-type-specific effects were observed. The strongest GO term for DEGs identified, extracellular matrix (ECM) organization, was shared between neurons, astrocytes, and microglia. This could reflect a more basic requirement for SORL1’s transport function or its function as a lipoprotein receptor for ECM maintenance. For example, many substrates involved in ECM integrity are dependent on the retromer complex for cell surface recycling (Sharma *et al*., 2020, Dong *et al*., 2013), and another lipoprotein receptor, LRP1, modulates ECM function in the CNS by both directly affecting proteins such as type III collagen (Salicioni *et al*., 2002, Gonias, Wu and Salicioni, 2004, Gaultier *et al*., 2006), and indirectly by regulating downstream signaling cascade of various scaffolding and adaptor proteins involved in ECM remodeling (Gaultier *et al*., 2010).

Most intriguing to us was the finding that SORL1 plays a neuron-specific role in the regulation of APOE and CLU levels. This finding is of particular interest given that in addition to SORL1, APOE and CLU variants also are strong genetic risk factors for AD. Little is known about the potential role of SORL1 as a lipoprotein receptor, and the regulation of APOE and CLU levels through SORL1 may be related to this role. In 2004, Scherzer *et al*. first showed reduced SORL1 expression in the AD brain and speculated that this was connected to its role as a neuronal APOE receptor (Scherzer *et al*., 2004). Findings of reduced SORL1 in the AD brain have since been replicated (Dodson *et al*., 2006, Sager *et al*., 2007), but there have been few studies since then studying the connection between SORL1 and APOE. These few studies used exogenously overexpressed SORL1 and APOE to identify their ability to interact, or the ability of SORL1 to mediate uptake of APOE in heterologous immortalized cell lines (Taira *et al*., 2001, Yajima *et al*., 2015). Here, with SORL1 knock out, APOE levels are reduced at both the protein and RNA levels in neurons but not in astrocytes or microglia. Unlike the elevation in Aβ and p-tau phenotypes observed in SORL1 null neurons, the neuron-specific reduction in APOE was not rescued by modulating retromer trafficking and autophagy.

In addition to protein-level changes, APOE RNA levels were reduced in SORL1 knock out iNs in two genetic backgrounds, and APOE RNA levels also were correlated with SORL1 expression across the ROSMAP iNs, suggesting that reductions in APOE protein levels were mediated at least in part at the level of transcription or RNA stability (Fig. 4B). Previous studies have examined transcriptional regulation of APOE in macrophages, hepatocytes, adipocytes, and astrocytes, but little is known about transcriptional regulation of APOE in neurons. Intriguingly, multiple pathway analyses performed herein indicated that TGF-β-mediated SMAD signaling was affected by SORL1 modulation. Components of this pathway were found to be differentially expressed at both the RNA and protein level with SORL1 KO (Fig. 1, Fig. 3I), and strongly correlated with SORL1 RNA and protein levels in the ROSMAP iN cohort (Fig. 5D). TGF-β has been shown to increase APOE expression in macrophages and monocytes (Singh and Ramji, 2006). Conversely, here we see that TGF-β treatment of iPSC derived neurons decreases APOE expression while SMAD inhibition increases APOE levels. SMAD-mediated regulation of transcription is highly complex and cell-type-specific effects on induction and suppression of transcription at the same genomic locus across different cell types have been reported (Derynck and Zhang, 2003, Luo, 2017), which may contribute to the neuron-specific effects on APOE RNA levels observed here. Indeed, ATAC-seq of WT cells revealed unique open chromatin sites at the APOE locus in differentiated neurons where SMADs may be recruiting additional factors that mediated APOE transcription. In this context, in iNs, SMADs appear to be acting to actively repress APOE transcription in neuronal cells, as TGF-β treatment of wild type neurons repressed APOE transcription while inhibition of SMAD signaling elevated APOE RNA levels (Fig. 7I). Of note, genes associated with SMAD signaling were down regulated at the RNA level with SORL1 knock out (Fig. 2I) which may reflect a negative feedback mechanism in response to elevated SMAD protein levels in the KO (Supp. Fig. 8G-I).

As described above, a large amount of genetic evidence supports a role for SORL1 in AD pathogenesis. While some rare truncation variants have been identified in early-onset AD individuals (Grear *et al*., 2009, Campion *et al*., 2019), there has been only one case of an individual with homozygous loss-of-function variants (Le Guennec *et al*. 2018). Thus, while the analyses of homozygous KO cells are helpful in understanding SORL1’s function, to more closely examine its contribution to AD risk, it was especially important to extend our analyses to probe the potential consequences of natural variation in SORL1 levels. Previous studies have shown that populations of individuals with AD and MCI have lower SORL1 levels in the brain than NCI populations, but little is known about the cause-and-effect relationships relevant to this finding. One study of iPSC-derived neurons differentiated from individuals with SORL1 risk haplotype (R) or protective haplotype (P) revealed that haplotype was not associated with the basal expression level of SORL1 mRNA (Young *et al*., 2015). Similarly, while our RNA and protein measurements from neurons and astrocytes showed a natural variation of SORL1 levels, there were no significant associations between SORL1 haplotypes and its expression levels. We speculate that a complex relationship between additional genetic variants at other loci interact with SORL1 haplotypes mediates its expression levels across individuals. Additional studies are warranted to define these genetic interactions, and a reductionist system such as the one described herein may be necessary to separate the genetic contribution to SORL1 expression from non-genetic environmental and age-related factors.

The roles of astrocytic and microglial APOE in neurodegeneration and AD neuropathology have been deeply investigated by many groups both *in vitro* and *in vivo*, but much less is known about the consequences of neuronal APOE expression. Questions remain regarding the implications of SORL1-mediated regulation of neuronal APOE. Neuronal APOE4 has been reported to cause an impairment of synaptic function (Lin *et al*., 2018), an increase in phosphorylation of tau and Aβ levels (Wang *et al*., 2018), and to accumulate in endosomes and impair endolysosomal trafficking (Xian *et al*., 2018). A recent study suggests that neuronal APOE may have a role in regulating inflammation (Zalocusky *et al*., 2021). In that study, human brain snRNAseq data revealed that neurons with low APOE expression have higher levels of MHC-I. Here, we probed that same human brain dataset and validated our *in vitro* findings of APOE and CLU association with SORL1 in excitatory neurons (Fig. 6). While SORL1 loss-of-function variants lead to AD, the reduction in APOE resulting from lower levels of SORL1 may represent a lowering of a protective mechanism in neurons, especially in APOE4 carriers. Data from the Herz lab and others suggest that APOE4 may fail to dissociate from its receptors within acidic endosomes due to its elevated isoelectric point relative to other APOE isoforms, and that the subsequent accumulation of APOE may underly the endosomal swelling in neurons observed in the AD brain (Nuriel *et al*. 2017, Xian *et al*., 2018). Aligned with this possibility, the iPSC line used to generate neurons that has an APOE ε3/ε4 haplotype had twice as many DEGs with SORL1 KO than the ε3/ε3 line (Supp. Table 1). Future studies are warranted to interrogate the consequences of SORL1 reduction in neurons of different APOE haplotypes in a genetically controlled manner.

### Limitations of study

The iPSC systems employed herein are reductionist systems that allow us to study the contribution of genetic variation of purified cell populations of defined fates. Here, we chose to study specific cell types in isolation to determine the cell autonomous roles of SORL1. However, using this simplified system we do not capture cell nonautonomous interactions that are certainly contributing to disease mechanisms. In addition, other factors that play an important role in AD such as aging are not captured. Lastly, while the differentiation protocols are continuously improving, iPSC-derived cell types cannot fully recapitulate the expression profiles of cells present in vivo in the brain. Despite these limitations, iPSC systems provide a well-controlled manipulatable system that allows us to identify cell biological processes affected by modulation of specific genes of interest, in this case SORL1. Hypotheses generated by iPSC studies can then be supported or challenged by data from complementary analyses of brain tissue. Here, our findings of a role for SORL1 in regulating levels of the apolipoproteins APOE and CLU in neurons are supported by data from human brain tissue and genetic studies of AD.

## Acknowledgements

We thank the NeuroTechnology Studio at Brigham and Women’s Hospital for providing Zeiss LSM710 Confocal instrument access and consultation on data acquisition and data analysis; iPSC NeuroHub at Brigham and Women’s Hospital for technical assistance with iN differentiation; Charles Jennings, Scott Small, and Gregory Petsko for manuscript feedback; Thomas Schwarz, Dennis Selkoe, and Francisco Quintana for their Dissertation Advisory Committee guidance, and to members of the Young-Pearse lab for their critical reading and manuscript feedback. This work was supported by NIH grants F31AG063399, U01AG072572, U01AG061356, RF1NS117446 and R01AG055909.

## Author Contributions

Conceptualization, H.L. and T.L.Y.-P.; Methodology, H.L., R.V.P., V.C., H.K., V.M., and T.L.Y.-P.; Software, R.V.P., H.K., V.M., and P.L.D.J.; Validation, H.L., A.J.A., Y.-C.H., Z.M.A., S.G., C.P., and D.M.D.; Formal Analysis, H.L., R.V.P., C.R.B., V.M., and T.L.Y.-P.; Investigation, H.L., A.J.A., Y.-C.H., Z.M.A., S.G., C.P., and D.M.D.; Resources, A.K., J.E.Y., D.A.B., P.L.D.J., and N.T.S.; Data Curation, R.V.P., D.A.B., P.L.D.J., and T.L.Y.-P.; Writing Original Draft, H.L. and T.L.Y.-P.; Writing – Review and Editing, A.J.A., R.V.P., Y.-C.H., Z.M.A., C.R.B., A.K., C.P., S.G., D.M.D., N.T.S., D.A.B., P.L.D.J., V.M., J.E.Y., and T.L.Y.P.; Visualization, H.L., R.V.P., C.R.B., H.K., V.M., and T.L.Y.-P.; Supervision, N.T.S., J.E.Y., V.M., D.A.B., P.L.D.J., and T.L.Y.-P.; Project Administration, T.L.Y.-P.; Funding Acquisition, D.A.B., P.L.D.J., and T.L.Y.-P.

## Figure Legends

## Materials and Methods

### Induced Pluripotent Stem Cell lines

iPSCs were generated from cryopreserved peripheral blood mononuclear cell (PBMC) samples from autopsied participants from either the Religious Order Study (ROS) or Rush Memory and Aging Project (MAP). ROS and MAP are cohort studies of aging and dementia. Each participant signed an informed consent, Anatomic Gift Act, and Repository Consent to allow their data to be repurposed, following approval by an Institutional Review Board of Rush University Medical Center. iPSCs were generated using Sendai reprogramming method as previously published (Lagomarsino *et al*., 2021). iPSCs undergo a rigorous quality procedure that includes a sterility check, mycoplasma testing, karyotyping, and pluripotency assays performed by the New York Stem Cell Foundation (NYSCF). iPSC cell line used to generate line 1 SORL1 KO cells was previously described and published (Young *et al*., 2015). iPSC cell line used to generate line 2 has been previously described (Muratore *et al*., 2014, Muratore *et al*., 2017), generated in collaboration with the Harvard Stem Cell Institute. The parental iPSC line (HVRDi002-A) harbored a APPV717I mutation, which was corrected using CRISPR/Cas technology (Muratore *et al*., 2017). APOE ε3/ε3 (cat#iPS26), APOE ε4/ε4 (cat#iPS16), and APOE KO (cat#iPS36) iPSCs were from obtained from Alstem. All three lines are isogenic, sharing the same genetic background, and APOE ε4/ε4 (cat#iPS16) was used as a parental line to edit APOE sequence. iPSCs were maintained using StemFlex Medium (Thermo Fisher Scientific). All cell lines were routinely tested for mycoplasma using PCR kit (MP0035-1KT) and STR profiling to prevent potential contamination or alteration to the cell lines.

### CRISPR/Cas9 to generate SORL1 KO iPSCs

Description and characterization of SORL1 WT/KO iPSCs in line 1 have been previously published (Knupp *et al*., 2020). gRNA ‘ATTGAACGACATGAACCCTC’ was used to target exon 6 at the VPS10p domain. To generate SORL1 WT/KO iPSCs in line 2, gRNA was chosen from Genescript’s website, which designs the gRNAs based on the algorithms by the Broad Institute. The gRNA sequence ‘ATGTTCCTGAATCATGATCC’ was used to target exon 5 at the VPS10p domain. IDT CRISPR Cas9 guide RNA design checker was used to check for any potential off-target effects.

### iPSC-derived neuron differentiation

iPSC-derived neurons (iNs) were differentiated following a previously published paper (Zhang *et al*., 2013) with minor modifications (Muratore *et al*., 2017, Lagomarsino *et al*., 2021). iPSCs were plated at a density of 95k cells/cm^2^ on plates coated with growth factor reduced Matrigel one day prior to virus transduction (Corning #354230). Then, iPSCs were transduced with three lentiviruses – pTet-O-NGN2-puro (Addgene plasmid #52047, a gift from Marius Wernig), Tet-O-FUW-EGFP (Addgene plasmid #30130, a gift from Marius Wernig), and FUdeltaGW-rtTA (Addgene plasmid #19780, a gift from Konrad Hochedlinger). The cells were then replated at 200,000 cells/cm^2^ using StemFlex Medium (Thermo Fisher Scientific) and ROCK inhibitor (10uM) (D0). The media was changed to KSR media (D1), 1:1 of KSR and N2B media (D2) and N2B media (D3). On day 4, cells were dissociated using Accutase, and plated at 50,000 cells/cm^2^ using iN D4 media (NBM media + 1:50 B27 + BDNF, GDNF, CNTF (10 ng/ml, Peprotech). Doxycycline (2ug/ml, Sigma) was added from D1 to the end of the differentiation, and puromycin (5 ug/ml, Gibco) was added from D2 to the end of the differentiation. On D3, B27 supplement (1:100) (Life Technologies) was added. From D4 to the end of differentiation D21, cells were cultured iN D4 media and fed every 2-3 days.

Induced neuron protocol media:

- <u>KSR media</u>: Knockout DMEM, 15% KOSR, 1x MEM-NEAA, 55 uM beta-mercaptoethanol, 1x GlutaMAX (Life Technologies).
- <u>N2B media</u>: DMEM/F12, 1x GlutaMAX (Life Technologies), 1x N2 supplement B (Stemcell Technologies), 0.3% dextrose (D-(+)-glucose, Sigma).
- <u>NBM media</u>: Neurobasal medium, 0.5x MEM-NEAA, 1x GlutaMAX (Life Technologies), 0.3% dextrose (D-(+)-glucose, Sigma).

### iPSC-derived astrocyte differentiation

iPSC-derived astrocytes (iAs) were differentiated following a previously published paper (Canals *et al*., 2018) with minor modifications (Lagomarsino *et al*., 2021). iPSCs were plated at 95k cells/cm^2^ on growth factor reduced matrigel (Corning #354230) coated plates prior to virus transduction. Then, iPSCs were transduced with three lentiviruses – Tet-O-SOX9-puro (Addgene plasmid #117269), Tet-O-NFIB-hygro (Addgene plasmid #117271), and FUdeltaGW-rtTA (Addgene plasmid #19780). The cells were then replated at 200,000 cells/cm^2^ using StemFlex Medium (Thermo Fisher Scientific) and ROCK inhibitor (10uM) (D0). The media was changed daily with Expansion Media (EM) from D1 to 3, and gradually switched from EM to FGF media from D4 to D7. On day 8, cells were dissociated using Accutase, and plated at 84,000 cells/cm^2^ using FGF media. Doxycycline (2.5ug/ml, Sigma) was added from D1 to the end of the differentiation, puromycin (1.25ug/ml, Gibco) was added on D3 of the differentiation, and hygromycin (100ug/ml, InvivoGen # ant-hg-1) was added from D4-D6 of the differentiation. From D8 to the end of differentiation D21, cells were cultured with maturation media and fed every 2-3 days.

Induced astrocyte protocol media:

- <u>Expansion Media</u>: DMEM/F12 (Thermo Fisher Scientific), 10% FBS, 1% N2 Supplement (Stemcell Technologies), 1% GlutaMAX (Life Technologies)
- <u>FGF Media</u>: Neurobasal media, 2% B27, 1% NEAA, 1% GlutaMAX, 1% FBS, 8ng/ml FGF, 5ng/ml CNTF, 10ng/ml BMP4
- <u>Maturation Media</u>: 1:1 DMEM/F12 and neurobasal media, 1% N2, 1% GlutaMAX, 1% Sodium Pyruvate, 5ug/ml N-1% N2, 1% GlutaMAX, 1% Sodium Pyruvate, 5ug/ml N-N-acetyl cysteine, 5ng/ml heparin-binding EGF-like GF, 10ng/ml CNTF, 10ng/ml BMP4, 500ug/ml dbcAMP

### iPSC-derived microglia-like cells differentiation

iPSC-derived microglia-like cells (iMGLs) were differentiated following a previously published paper (Abud *et al*., 2017, McQuade et al.. 2018) with minor modifications (Lagomarsino *et al*., 2021, Chou et al. in submission). iPSCs were plated on growth factor reduced matrigel (Corning #354230) using StemFlex Medium (Thermo Fisher Scientific) and ROCK inhibitor (10uM). From D0 to D12, StemDiff Hematopoietic Kit (Stemcell Technologies) was used to generated hematopoietic precursor cells (HPCs). At D12, cells were replated at 100,000 cells per 35mm well in iMGL media supplemented with 3 cytokines (IL-34 (100ng/mL, Peprotech), TGF-β1 (50ng/mL, Militenyi Biotech), M-CSF (25ng/mL, Thermo Fisher Scientific)). From D12 to D24, iMGL media with freshly added cytokines were added to the culture every other day. On D24, cells were replated at 100,000 cells per 15.6mm well with 1:1 mixture of old media and fresh iMGL media with 3 cytokines. From D24 to D37, iMGL media with freshly added 3 cytokines were added to the culture every other day. On D37, cells are resuspended in iMGL media with five cytokines, supplemented every other day until D40.

iPSC-derived microglia-like cells protocol media:

- iMGL media: DMEM/F12, 2X insulin-transferrin-selenite, 2X B27, 0.5X N2, 1X GlutaMAX, 1X non-essential amino acids, 400μM monothioglycerol, 5 μg/mL insulin
- 3 cytokines: 100 ng/mL IL-34 (Peprotech), 50 ng/mL TGF-β1 (Millitenyi Biotech), and 25 ng/mL M-CSF (ThermoFisher Scientific)
- 5 cytokines: 100 ng/mL IL-34, 50 ng/mL TGF-β1, 25 ng/mL M-CSF, 100 ng/mL, CD200 (Novoprotein) and 100 ng/mL CX3CL1 (Peprotech)

### iPSC-derived brain microvascular endothelial cells differentiation

iPSC-derived brain microvascular endothelial cells (iBMVECs) were differentiated following previously published papers (Lippmann *et al*., 2012, Lippmann *et al*., 2015, Park *et al*., 2019) with minor modifications (Chou et al. in submission). iPSCs were plated on growth factor reduced matrigel (Corning #354230) using StemFlex Medium (Thermo Fisher Scientific) and ROCK inhibitor (10uM) at a density of 180,000 cells per 35mm well (D-2). From D0 to D4, the media was changed daily with E6 medium (ThermoFisher) using hypoxia condition (5%CO2-5%O2-N2). From D5, the media was changed to EC media supplemented with 20ng/ml FGF and 10uM retinoic acid. On D7, the media is replaced with new EC media supplemented with fresh FGF and retinoic acid. From D8-D14, the media was changed daily with EC medium until harvest.

- Endothelial media (EC): hESFM (Thermo Fisher, Cat#: 11111044), 50X B27 supplement (Thermo Fisher, Cat#: 17504044), and (0.5%) pen/strep (Thermo Fisher, Cat#: 15140163)

### Western Blotting

Cells were lysed with the RIPA lysis buffer (Thermo Fisher Scientific #89900) or in a buffer containing 1%NP40, 10mM EDTA, 150mM NaCl, 50mM Tris, with the protease inhibitor (cOmplete™ mini protease inhibitor, Roche) and phosphatase inhibitor (phosphoSTOP, Roche) added freshly before the lysis. Using BCA Protein Assay Kit (Pierce), protein concentrations were measured, and equal amount of protein were loaded onto 4-12% Bis-Tris NuPAGE gels (Life Technologies). Gels were then transferred to nitrocellulose membranes, blocked with LI-COR TBS blocking buffer for 1 hour, then probed with primary antibodies overnight. After washing three times in TBST (0.05% Tween in TBS), the secondary antibodies were added for 1hr rocking in RT, followed by three more washes in TBST. Blots were imaged using Odyssey Clx system (LI-COR) and quantified using ImageStudio (LICOR).

Antibodies for Immunocytochemistry and Western Blot

| Antigen | Host | ICC/WB | Dilution | Vendor | Catalog # |
| --- | --- | --- | --- | --- | --- |
| APOE | Goat | ICC and WB | 1:1000 | Sigma | 178479 |
| APP | Rabbit | WB | 1:1000 | Sigma | A8717 |
| P202, 205 AT8 (MAPT) | Mouse | WB | 1:250 | ThermoFisher | MN1020 |
| GAPDH | Mouse | WB | 1:10,000 | ProteinTech | 60004 |
| GFAP | Chicken | ICC | 1:1000 | Abcam | ab4672 |
| IBA1 | Goat | ICC | 1:500 | Abcam | Ab5076 |
| LC3 | Rabbit | WB | 1:1000 | MBL | PM036 |
| MAP2 | Chicken | ICC | 1:5000 | Abcam | ab5392 |
| SORL1 (LR11) | Mouse | WB | 1:1000 | BD Biosciences | 611860 |
| SYN1 | Rabbit | ICC | 1:1000 | Calbiochem | 574777 |
| TAU (MAPT) | Rabbit | WB | 1:2000 | Dako | A0024 |
| TJP1 | Mouse | ICC | 1:1000 | Invitrogen | 339100 |
| TUJ1 | Mouse | ICC | 1:1000 | Milipore | MAB1637 |
| VIMENTIN | Mouse | ICC | 1:500 | Milipore | CBL202 |

### qPCR

At iN d21, cells were harvested and RNA was purified using Purelink RNA Mini kit (Invitrogen). cDNA was generated using SuperScript II (Invitrogen). qPCR was performed using Power SYBR™ Green Master Mix and run on ViiA7 system (Applied Biosystems).

### Primers

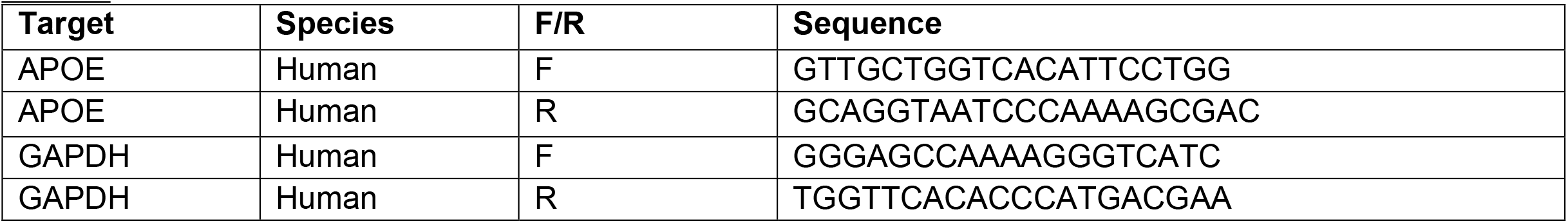

### ELISAs

48hr conditioned media was collected from d19 to d21 before harvest. Extracellular Aβ 38, 40, and 42 levels were measured using MSD V-PLEX Aβ Peptide Panel 1 (6E10) Kit (cat. # K15200G-1). Extracellular APOE and CLU were measured using MSD R-PLEX Human ApoE Assay (cat. # K1512IR-2) and MSD R-PLEX Human Clusterin Assay (cat. # K151YLR-2).

### Immunocytochemistry

At iN d21, iA d21, iMGL d40, and iBMVEC d14, cells were washed once with PBS, then was fixed in 4% paraformaldehyde solution for 15 minutes at RT. After completing the fixation, cells were incubated in blocking buffer (2% donkey serum with 0.1% Triton X-100) for 1 hour rocking at room temperature followed by overnight incubation in primary antibody at 4C. Then, cells are washed with PBS three times, incubated with secondary antibodies, and then washed with PBS three times. DAPI (1:1000) staining was performed during the second wash with PBS. Then, the cells were imaged using LSM710 confocal microscopy.

### Drug treatments

Prior to harvest, cells were fed with iN media containing the following drugs: 10uM or 20uM R33 (MedKoo Biosciences), 100mM Trehalose dihydrate (Sigma), and 5nM Chloroquine (Tocris) for 72hrs; 10ng/ml TGF-β (T7039) and 10uM SB-431542 (616464) for 48hrs (western blot) and 24hrs (qPCR). The vehicles were 1:1 mix of water and DMSO for R33, ethanol for SB-431542, and water for chloroquine and TGF-β. Trehalose was prepared directly in iN media. For trehalose and chloroquine, cells were treated one more time on D20 before final harvest on d21.

### Lentiviral transduction (shRNA)

Lentivirus expressing pLKO.1 vector was used to generate two APOE shRNA constructs. TRCN0000371278, referred to ‘APOE1’ in this study and TRCN0000377711, referred to ‘APOE2’ in this study were obtained from Sigma. iNs were transduced at D17 with either empty virus, APOE1, or APOE2, at MOI of 3. Following 18hrs of incubation, the media was replaced with fresh media and incubated for additional 72hr. At d21, media and cells were collected for further analysis.

### Bulk RNA sequencing

High quality total RNA (eRIN > 9.0) was harvested from differentiated iNs, iAs, iMGLs, iBMVECs, or from undifferentiated iPSCs. Libraries were generated and sequenced to a depth of 30 million read pairs using the GeneWiz next-gen sequencing service. RNAseq reads were quality trimmed to remove end base called with phred33 scores below 25 and eliminate resulting reads that are shorter than 30 bases. Trimmed reads are quantified using the Kalliso (v 0.43.1) pseudoalignment algorithm with 50 bootstraps (Bray *et al*., 2016)

### Genomic data analyses

TPM Normalized expression matrices and differential expression statistics were generated using the Sleuth (v 0.30) package in R (v 4.0.3). Expression differences were tested using a Wald test after controlling for differentiation batch. Significant hits were defined by q value (FDR) < .05 and a b value > .5 Gene Ontology enrichments were tested using the clusterProfiler package (v 3.18.1) (Wu *et al*., 2021) in R. Enrichment dot plots were generated using ggplot2 (v 3.3.5) (Wickham 2016) while the gene concept network analyses were generated using the enrichplot (v 1.12.2) (Yu 2022) package. Correlations were analyzed using custom R scripts and base correlation functions.

### Protein Digestion

Each cell pellet was individually homogenized in 300 uL of urea lysis buffer (8 M urea in 10 mM Tris, 100 mM NaH2PO4 buffer, pH=8.5), including 5 uL (100x stock) HALT protease and phosphatase inhibitor cocktail (Pierce). All homogenization was performed using a Bullet Blender (Next Advance) according to manufacturer protocols. Briefly, each tissue piece was added to Urea lysis buffer in a 1.5 mL Rino tube (Next Advance) harboring 750 mg stainless steel beads (0.9-2 mm in diameter) and blended twice for 5 minute intervals in the cold room (4°C). Protein supernatants were transferred to 1.5 mL Eppendorf tubes and sonicated (Sonic Dismembrator, Fisher Scientific) 3 times for 5 s with 15 s intervals of rest at 30% amplitude to disrupt nucleic acids and subsequently vortexed. Protein concentration was determined by the bicinchoninic acid (BCA) method, and samples were frozen in aliquots at −80°C. Protein homogenates (50ug) treated with 1 mM dithiothreitol (DTT) at 25°C for 30 minutes, followed by 5 mM iodoacetimide (IAA) at 25°C for 30 minutes in the dark. Protein mixture was digested overnight with 1:100 (w/w) lysyl endopeptidase (Wako) at room temperature. The samples were then diluted with 50 mM NH4HCO3 to a final concentration of less than 2M urea and then and further digested overnight with 1:50 (w/w) trypsin (Promega) at 25°C. Resulting peptides were desalted with a Sep-Pak C18 column (Waters) and dried under vacuum.

### Tandem Mass Tag (TMT) Labeling

Peptides were reconstituted in 100ul of 100mM triethyl ammonium bicarbonate (TEAB) and labeling performed as previously described (1, 2) using TMTPro isobaric tags (Thermofisher Scientific, A44520). Briefly, the TMT labeling reagents were equilibrated to room temperature, and anhydrous ACN (200 μL) was added to each reagent channel. Each channel was gently vortexed for 5 min, and then 20 μL from each TMT channel was transferred to the peptide solutions and allowed to incubate for 1 h at room temperature. The reaction was quenched with 5% (vol/vol) hydroxylamine (5 μl) (Pierce). All 16 channels were then combined and dried by SpeedVac (LabConco) to approximately 100 μL and diluted with 1 mL of 0.1% (vol/vol) TFA, then acidified to a final concentration of 1% (vol/vol) FA and 0.1% (vol/vol) TFA. Peptides were desalted with a 60 mg HLB plate (Waters). The eluates were then dried to completeness. High pH fractionation was performed essentially as described (3) with slight modification. Dried samples were re-suspended in high pH loading buffer (0.07% vol/vol NH4OH, 0.045% vol/vol FA, 2% vol/vol ACN) and loaded onto a Water’s BEH (2.1mm × 150 mm with 1.7 μm beads). An Thermo Vanquish UPLC system was used to carry out the fractionation. Solvent A consisted of 0.0175% (vol/vol) NH4OH, 0.01125% (vol/vol) FA, and 2% (vol/vol) ACN; solvent B consisted of 0.0175% (vol/vol) NH4OH, 0.01125% (vol/vol) FA, and 90% (vol/vol) ACN. The sample elution was performed over a 25 min gradient with a flow rate of 0.6 mL/min with a gradient from 0 to 50% B. A total of 96 individual equal volume fractions were collected across the gradient and dried to completeness using a vacuum centrifugation.

### Liquid chromatography Coupled to Tandem Mass Spectrometry (LC-MS/MS)

All samples were analyzed on the Evosep One system using an in-house packed 15 cm, 75 μm i.d. capillary column with 1.9 μm Reprosil-Pur C18 beads (Dr. Maisch, Ammerbuch, Germany) using the pre-programmed 21 min gradient (60 samples per day) essentially as described (4). Mass spectrometry was performed with a high-field asymmetric waveform ion mobility spectrometry (FAIMS) Pro equipped Orbitrap Eclipse (Thermo) in positive ion mode using data-dependent acquisition with 2 second top speed cycles. Each cycle consisted of one full MS scan followed by as many MS/MS events that could fit within the given 2 second cycle time limit. MS scans were collected at a resolution of 120,000 (410-1600 m/z range, 4×10^5 AGC, 50 ms maximum ion injection time, FAIMS compensation voltage of -45). All higher energy collision-induced dissociation (HCD) MS/MS spectra were acquired at a resolution of 30,000 (0.7 m/z isolation width, 35% collision energy, 1.25×10^5 AGC target, 54 ms maximum ion time, TurboTMT on). Dynamic exclusion was set to exclude previously sequenced peaks for 20 seconds within a 10-ppm isolation window.

### Database Searching and Protein Quantification

All raw files were searched using Thermo’s Proteome Discoverer suite (version 2.4.1.15) with Sequest HT. The spectra were searched against a human uniprot database downloaded August 2020 (86395 target sequences). Search parameters included 10ppm precursor mass window, 0.05 Da product mass window, dynamic modifications methione (+15.995 Da), deamidated asparagine and glutamine (+0.984 Da), phosphorylated serine, threonine and tyrosine (+79.966 Da), and static modifications for carbamidomethyl cysteines (+57.021 Da) and N-terminal and Lysine-tagged TMT (+304.207 Da). Percolator was used filter PSMs to 0.1%. Peptides were group using strict parsimony and only razor and unique peptides were used for protein level quantitation. Reporter ions were quantified from MS2 scans using an integration tolerance of 20 ppm with the most confident centroid setting. Only unique and razor (i.e., parsimonious) peptides were considered for quantification. All RAW FILES are available on PRIDE

### snRNAseq

Raw counts data from the Mathys et al. (2019) study were converted to Counts Per Million (CPM) by dividing counts for each nucleus by the sum of counts for that nucleus and then multiplying by 10^6. From this normalized matrix, the following analyses were carried out:

1. Differential expression of APOE among SORL1+ versus SORL1-nuclei. For each major cell class, nuclei were separated into SORL1+ (with expression>0) and SORL1- (with expression=0) groups, and a Mann-Whitney test was performed (using the wilcox.test function in R) to determine differential expression of APOE. Raw p-values were adjusted with Bonferroni correction (p-value*number of tests performed). The fold change was calculated as the ratio of the log-transformed (log10(CPM+1)) mean expression of APOE in the SORL1+ group divided by the log-transformed mean expression of APOE in the SORL1-group. This same analysis was performed for differential expression of CLU between the two groups as well.
2. Significance of overlap in detection of APOE among SORL1+ nuclei. For each major cell class, a Fisher’s exact test was performed to determine whether the number of double-positive nuclei (APOE+/SORL1+) was greater or less than expected by chance. Odds ratio (with upper and lower bounds) were extracted from these results and graphed. Raw p-values were adjusted with Bonferroni correction (p-value*number of tests performed). This same analysis was performed to assess the significance of overlap in detection of CLU among SORL1+ nuclei.

### ATAC-seq

ATAC libraries were sequencing using Illumina NovaSeq generating paired-end reads of 50bp lengths. Reads were aligned against the GRCh38 reference genome using Bowtie2 vers. 2.3.5.1 in the very-sensitive-local mode (Langmead and Salzberg, 2012). Quality metrics were calculated using the Picard tools (see Supp. Table 7). Only proper pairs of reads were passed to Macs2 vers. 2.1.2 for peak calling with a q-value threshold of 0.05 (Zhang *et al*., 2008). Standardized coverage tracks for visualization were generated by the Bedtools suite vers. 2.30. The R package TFBSTools vers. 1.32.0 was used to detected putative SMAD binding sites in accessible regions at the APOE locus using the motifs MA0795_1, MA1153_1, MA1557_1, and MA1964_1 from the JASPAR 2022 database (Tan and Lenhard, 2016, Castro-Mondragon *et al*., 2022).

### Statistical analysis

All statistical tests were performed using GraphPad Prism 9 or R Studio. All data is shown as mean +/- SEM. Comparisons between two groups were using student’s t test, and comparisons between more than two groups were analyzed using one-way ANOVA followed by Tukey’s post-hoc test. Spearman correlation method (rank-order based) and Pearson correlation method (linear relationship) was used for correlation analyses.

## Supplemental Figure legends

**Supp. Figure 1.**
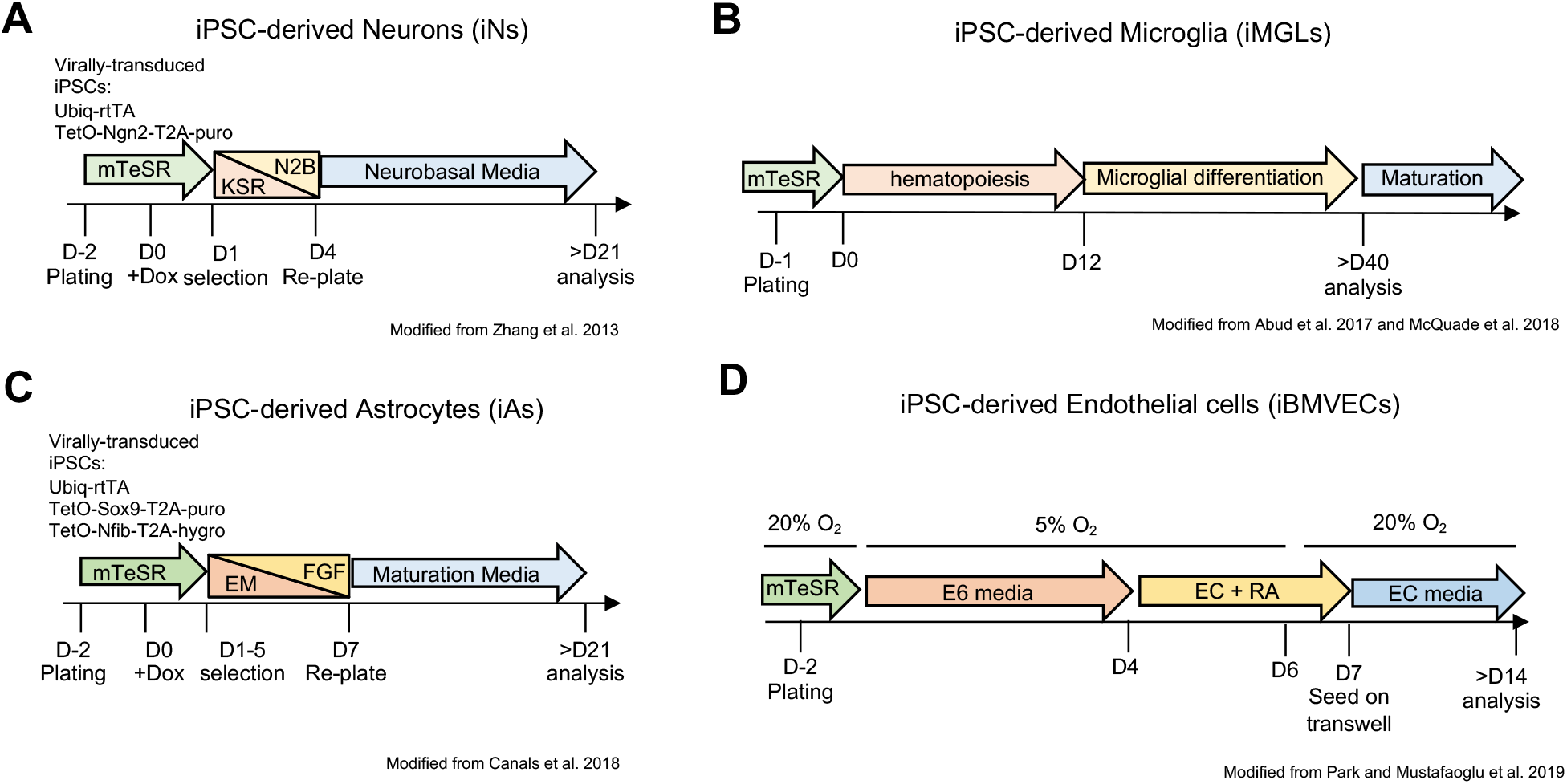
Overview of iPSC differentiation protocols. Schematic of (A) iN, (B) iMGL, (C) iA, and (D) iBMVEC differentiation protocol used in this study.

**Supp. Figure 2.**
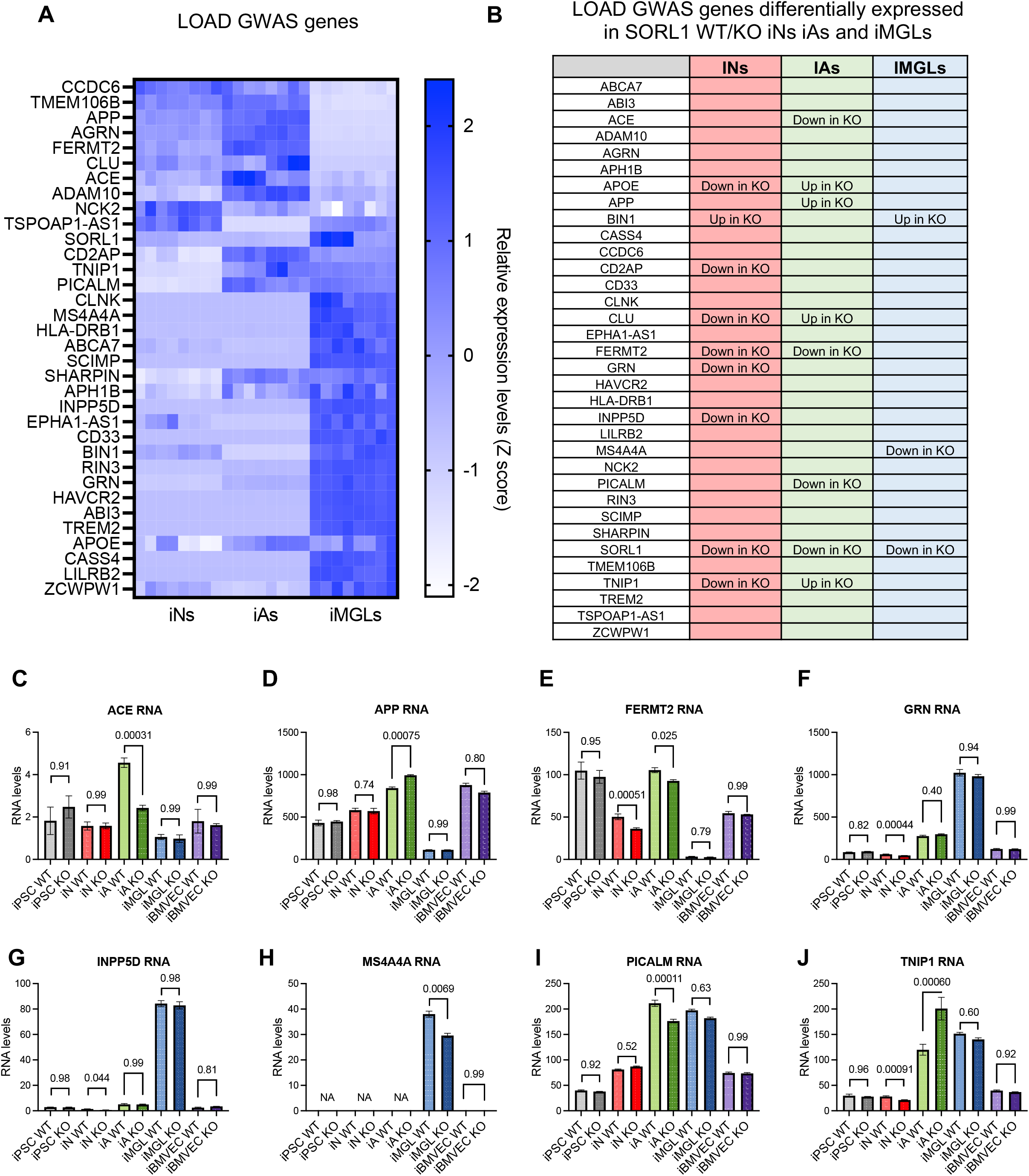
Expression of LOAD GWAS genes in SORL1 WT/KO iNs, iAs, iMGLs. (A) Heatmap of RNA expression levels of LOAD GWAS genes in iNs, iAs, and iMGLs. (B) Table of known LOAD GWAS genes differentially expressed in SORL1 WT/KO neurons, astrocytes, or microglia. (C-J) RNA levels of LOAD GWAS genes in 5 different cell types (Genes not listed in Figure 2).

**Supp. Figure 3.**
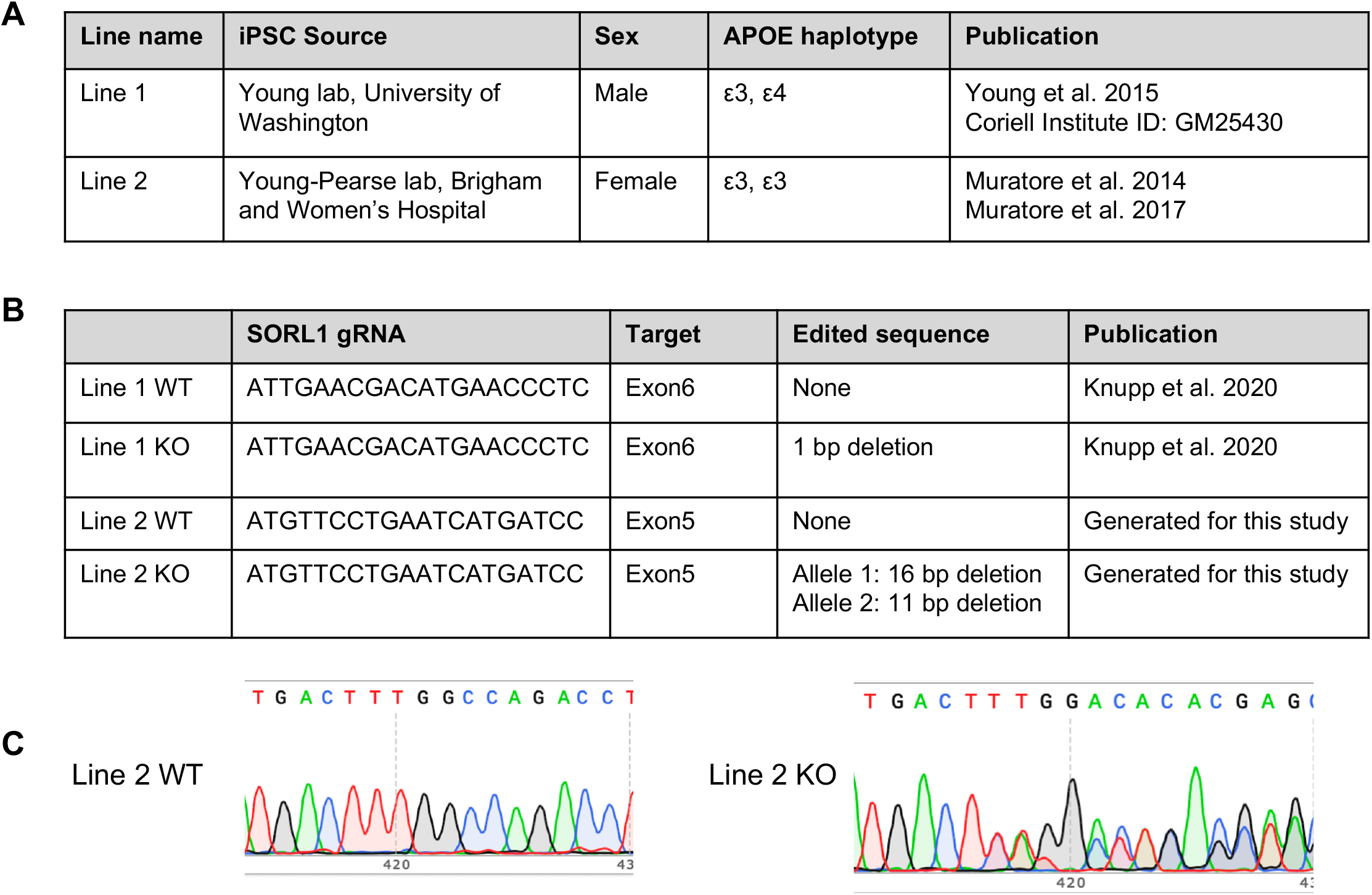
Generation of SORL1 KO iPSCs using CRISPR/Cas9. (A) Table showing the iPSC line source information. (B) Table with the information on the gRNAs used to target SORL1, and the edited sequence results for each clone used in this paper. (C) Sequence chromatogram of Line 2 WT and line 2 KO lines.

**Supp. Figure 4.**
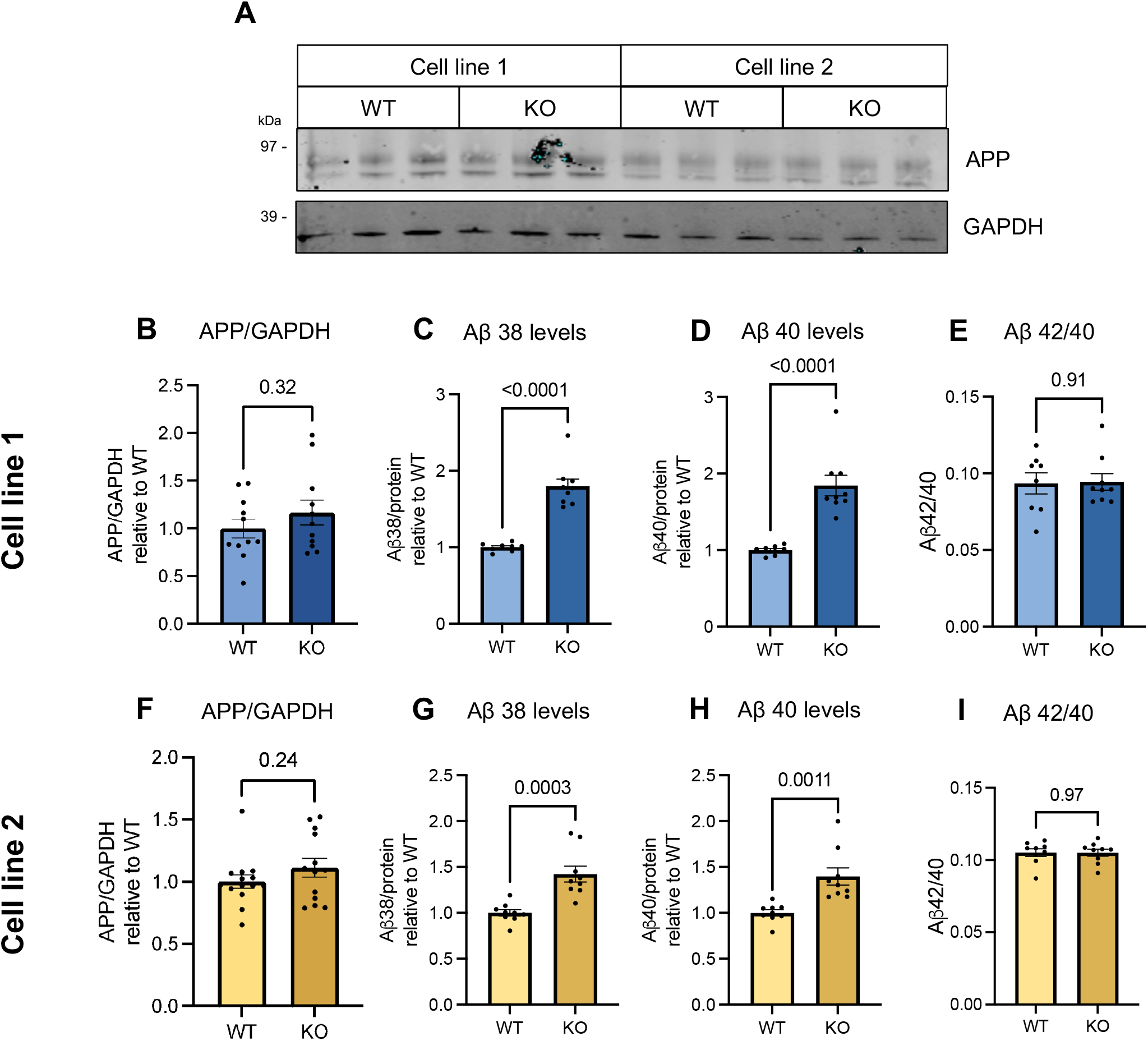
SORL1 KO neurons show increased Aβ levels without altering APP levels. (A) Representative western blot of APP protein expression in SORL1 WT and KO iNs in cell line 1 and 2. (B,F) Quantification of APP/GAPDH in iNs.(C,G) Aβ38 levels, (D,H) Aβ40 levels, and (E,I) Aβ 42/40 ratio in media generated from SORL1 WT and KO iNs in cell lines 1 and 2. Data show mean +/- SEM from three differentiations, n=3 per differentiation for each line. Values are normalized to WT for each pair. Unpaired student’s t test (two tailed).

**Supp. Figure 5.**
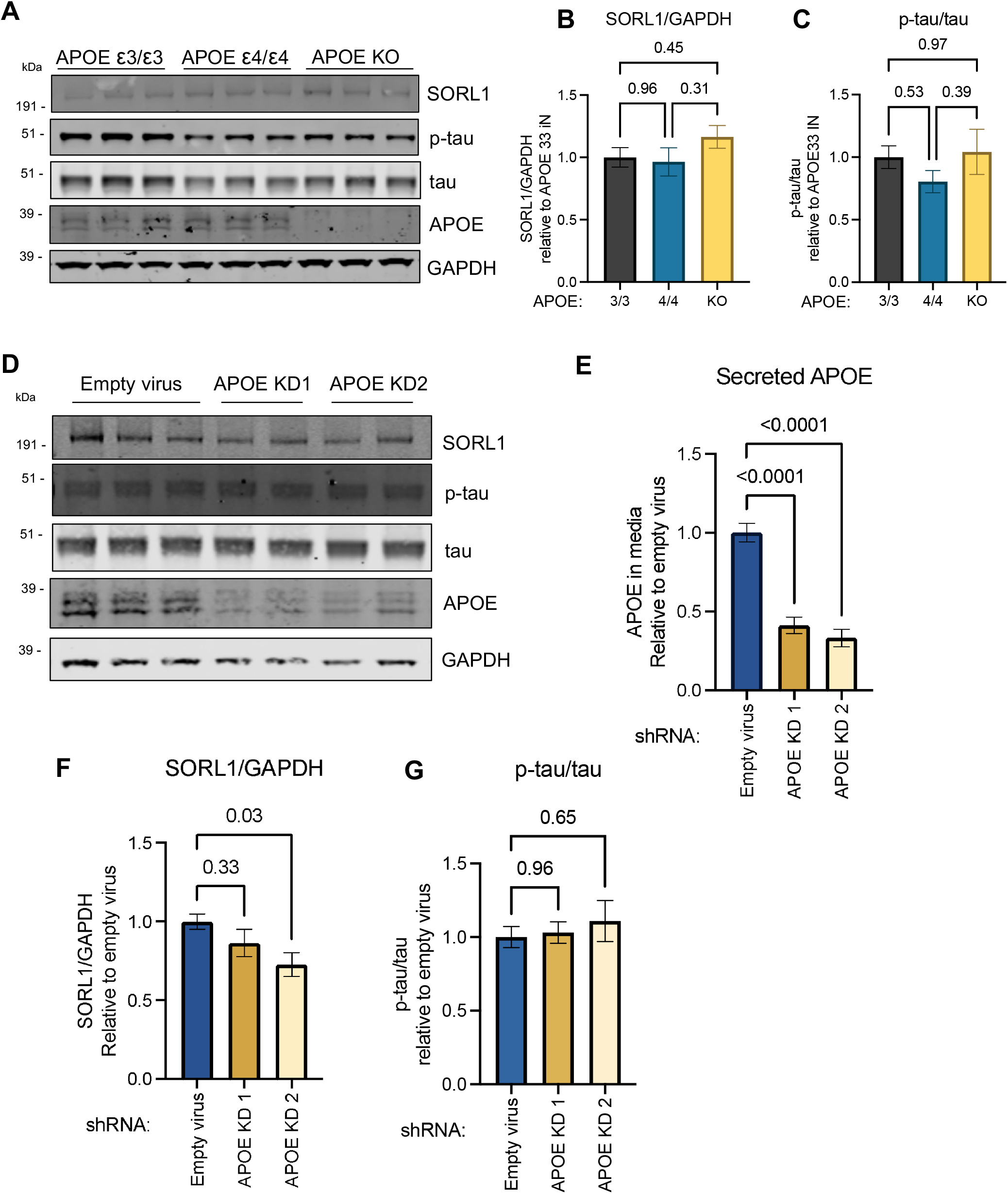
Loss of APOE does not affect SORL1 levels and p-tau levels in iNs. (A) Representative western blot of an isogenic set of iNs with APOE ε3/ε3, APOE ε4/ε4, and APOE KO genotypes. (B) Quantification of SORL1/GAPDH and (C) p-tau/tau. (D) Representative western blot of iNs that received APOE shRNA KD transduction. (E) Quantification of secreted APOE in the media, (F) SORL1/GAPDH, and (G) p-tau/tau. Data show mean +/- SEM from three differentiations, n=2-3 per differentiation for each line. Values are normalized to APOE ε3/ε3 iNs or iNs with empty virus. One-way ANOVA with Tukey’s multiple comparisons test.

**Supp. Figure 6.**
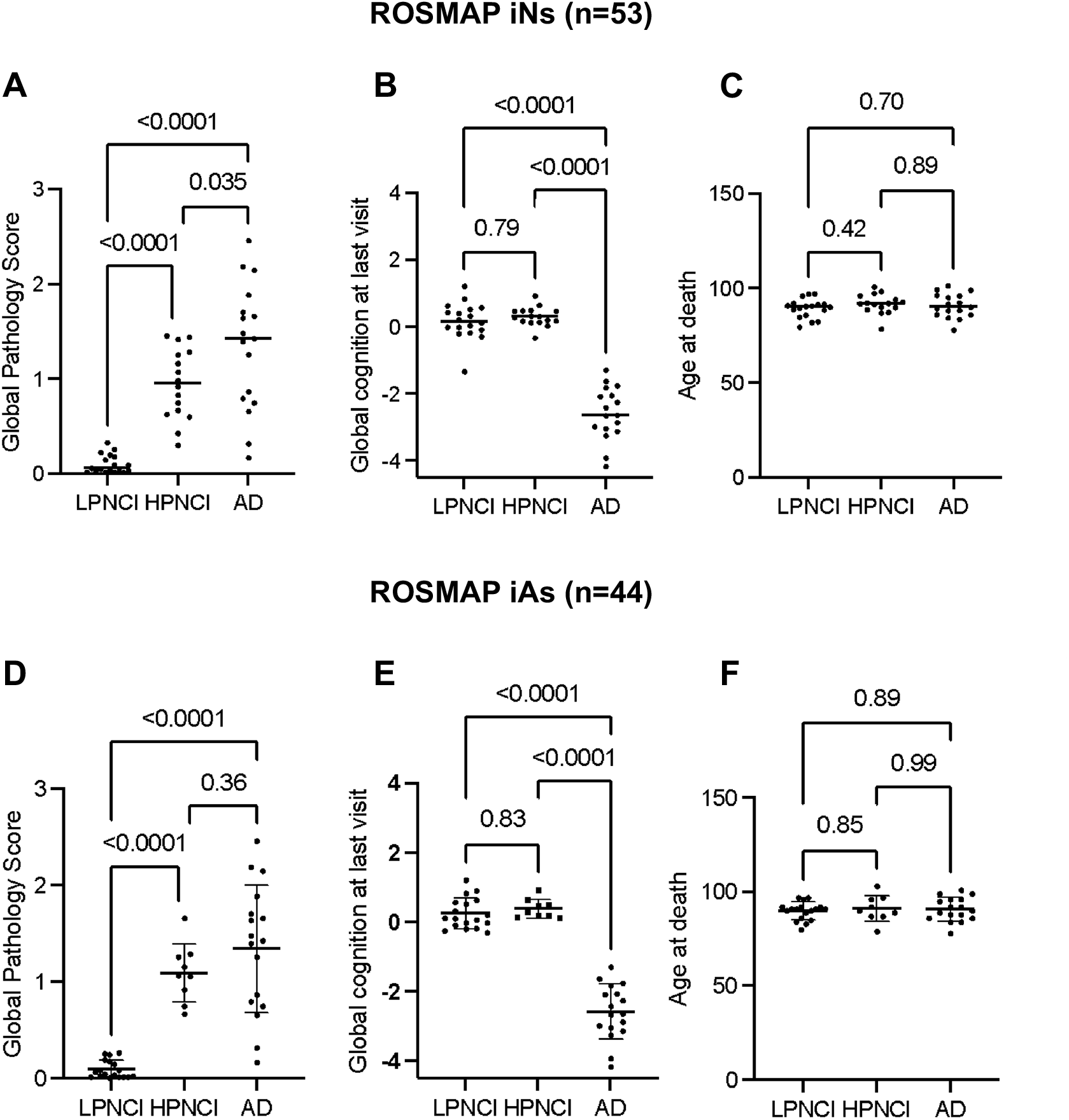
Overview of ROSMAP cohort participants. (A,D) Global Pathology Score (B,E) Cognition at Last Visit and (C,F) Age at Death of individuals from ROSMAP cohort used to generate iNs and iAs. Individuals in LPNCI show low pathology and no dementia, while individuals in HPNCI category show high pathology and no dementia.

**Supp. Figure 7.**
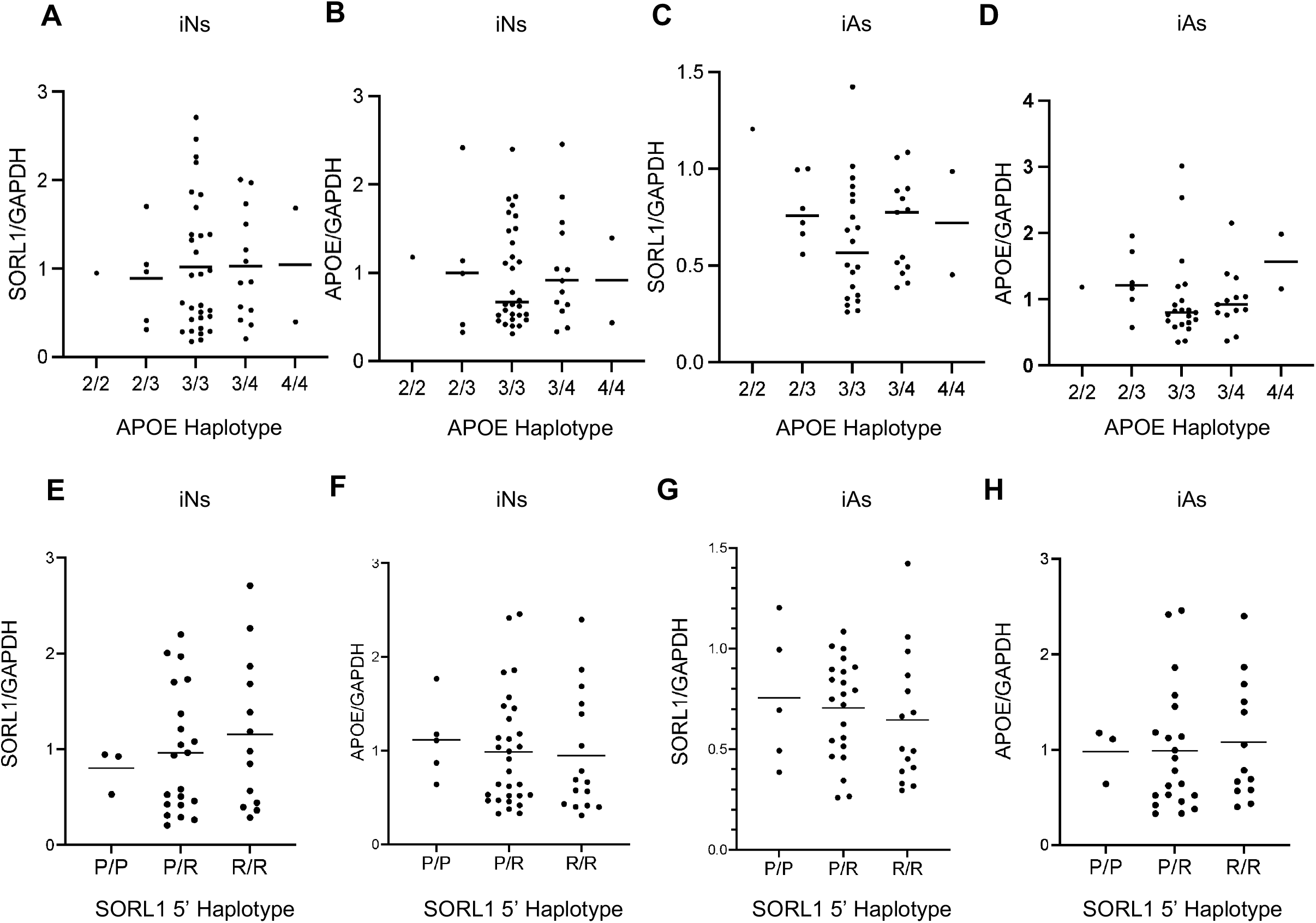
SORL1 and APOE expression in iNs and iAs based on SORL1 and APOE haplotype. (A,C) SORL1 protein levels and (B,D) APOE protein levels in iNs and iAs western blots from the ROSMAP cohort categorized based on APOE haplotype. (E,G) SORL1 protein levels and (F,H) APOE protein levels in iNs and iAs western blots from the ROSMAP cohort categorized based on SORL1 5’ haplotype. Data show mean +/- SEM. One-way ANOVA with Tukey’s multiple comparisons test.

**Supp. Figure 8.**
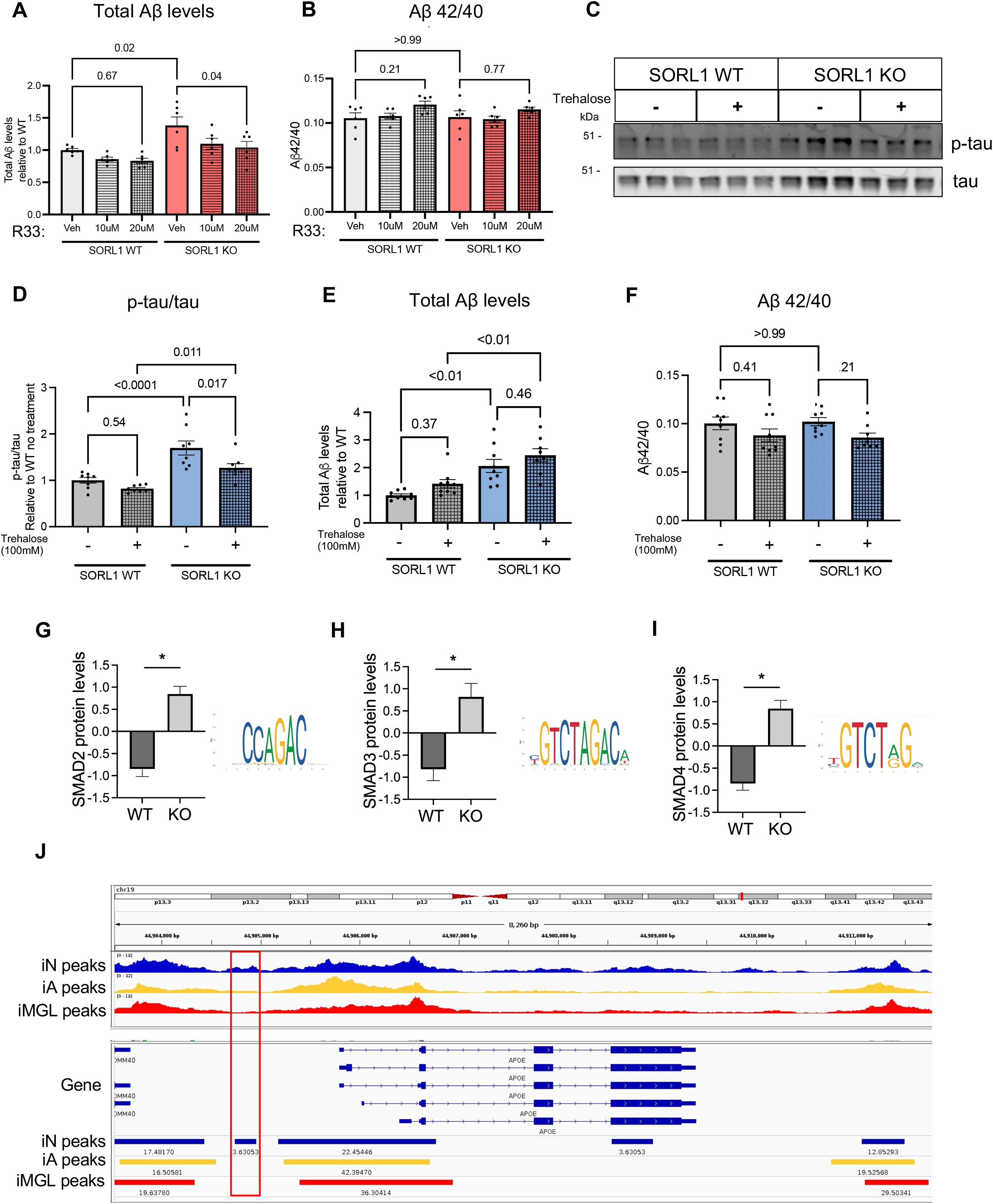
Interrogation of candidate pathways associated with SORL1 loss-of-function. (A) Total Aβ levels after 72hr treatment of d17 iNs with 10uM or 20uM R33. Elevated Aβ levels in SORL1 KO iNs are partially rescued with 20 uM R33. (B) Aβ42/40 is not altered with R33 treatment. (C) A representative western blot of SORL1 WT and KO iNs treated with trehalose (100mM) for 72 hours. (D) Quantification of p-tau/tau (p202/205). (E) Total Aβ levels in media of iNs after 72hr treatment with 100mM trehalose treatment. Elevated Aβ levels in SORL1 KO iNs are not rescued with trehalose. (F) Aβ42/40 is not altered with trehalose treatment. Data show mean +/- SEM from three differentiations, n=3 per differentiation for each line. Values are normalized to WT iNs treated with vehicle. One-way ANOVA with Tukey’s multiple comparisons test. (G-I) TMT-MS results of SORL1 WT and KO iNs show an elevation in protein levels of SMAD2, SMAD3, and SMAD4. DNA binding consensus sites are shown, acquired from the JASPAR database (Castro-Mondragon *et al*., 2022). (J) ATAC-seq was performed on iNs, iAs, and iMGLs differentiated in parallel from an iPSC line derived from an individual with AD (BR98). Shown is the Integrative Genomics Viewer (IGV, Broad Institute) image of coverage of mapped reads to the APOE locus for iNs, iAs, and iMGLs. Red box highlights region of open chromatin present in iNs that shows reduced accessibility in iAs and iMGLs.

## Supplemental Tables

**Supp. Table 1**. DEGs in SORL1 KO vs. WT iPSCs, iNs, iAs, iMGLs, and iBMVECs.

**Supp. Table 2**. Enriched GO pathways in SORL1 KO vs. WT iPSCs, iNs, iAs, iMGLs, and iBMVECs.

**Supp. Table 3**. Enriched GO pathways in SORL1 KO vs. WT iNs in both line 1 and line 2.

**Supp. Table 4**. Enriched GO pathways in SORL1 KO vs. WT neurons using DEGs from Mishra *et al*. 2022.

**Supp. Table 5**. Detection overlap and differential expression analysis of snRNAseq data from Mathys *et al*. 2019

**Supp. Table 6**. SMAD binding sites in the human APOE promoter corresponding to regions of open chromatin.

**Supp. Table 7**. Quality metrics used in the ATAC-seq analysis of the APOE locus.

